# Gene-dependent yeast cell death pathway requires AP-3 vesicle trafficking leading to vacuole membrane permeabilization

**DOI:** 10.1101/2021.08.02.454728

**Authors:** Zachary D. Stolp, Madhura Kulkarni, Yining Liu, Chengzhang Zhu, Alizay Jalisi, Si Lin, Arturo Casadevall, Kyle W. Cunningham, Fernando J. Pineda, Xinchen Teng, J. Marie Hardwick

## Abstract

Unicellular eukaryotes are suggested to undergo self-inflicted destruction. However, molecular details are sparse by comparison to the mechanisms of cell death known for human cells and animal models. Here we report a molecular pathway in *Saccharomyces cerevisiae* leading to vacuole/lysosome membrane permeabilization and cell death. Following exposure to heat-ramp conditions, a model of environmental stress, we observed that yeast cell death occurs over several hours, suggesting an ongoing molecular dying process. A genome-wide screen for death-promoting factors identified all subunits of the AP-3 adaptor complex. AP-3 promotes stress-induced cell death through its Arf1-GTPase-dependent vesicle trafficking function, which is required to transport and install proteins on the vacuole/lysosome membrane, including a death-promoting protein kinase Yck3. Time-lapse microscopy revealed a sequence of events where AP-3-dependent vacuole permeability occurs hours before the loss of plasma membrane integrity. An AP-3-dependent cell death pathway appears to be conserved in the human pathogen *Cryptococcus neoformans*.

## Introduction

Two long-standing conventions have challenged the existence of programmed or regulated cell death (PCD/RCD) mechanisms in unicellular eukaryotes. First, from observations of orderly patterns of cell death in developing animals (Lockshin and Williams, 1965), programmed death was thought to arise during the evolution of multicellular organisms. Second, for decades the prevailing evolution theory rejected the possibility of cell suicide as an adaptation (the ultimate altruistic behavior) in unicellular organisms because this conflicted with the concept of individual-level theory of selection (versus multi- or group-level selection) (Durand, 2020). Supporting both of these ideas, unicellular eukaryotes and bacteria lack key molecular players of classical apoptosis, the best-studied cell death pathway in mammals (BAX-inducible, caspase-3-mediated cell fragmentation and engulfment by neighboring cells) (Kuida et al., 1996; Lindsten et al., 2003; Nagata and Segawa, 2021). However, these conventions are not mutually exclusive with programmed death in unicellular organisms.

Accumulating evidence has prompted reappraisal of these conventions, including observations of cell suicide as a defense mechanism in bacteria (Iranzo et al., 2014). The emerging alternative concept is that programmed unicellular death predates and was required for the emergence of multicellular organisms, rather than the inverse (Koonin and Zhang, 2017). While there is little doubt that selfish genes drive evolution of cell death (Ramisetty et al., 2015), this does not preclude group level selection theory explanations that accommodate unicellular programmed cell death (Durand, 2020). Similarly, while classic apoptosis (caspase-3-mediated cell death) likely arose with metazoans where it plays a key role in ontogeny, its absence in unicellular organisms does not reject the concept of self-inflicted unicellular death, as the molecular details of cell death may be as diverse as the organisms themselves (Ameisen, 2002; Teng and Hardwick, 2015). However, new parallels between microorganisms and mammals have emerged for non-apoptotic programmed death mechanisms. For example, pore-forming, cell suicide-inducing gasdermin proteins were recently found in bacteria and multi-cellular/filamentous fungi (Clavé et al., 2021; Johnson et al., 2021). Despite low sequence similarity, they share striking structural and mechanistic similarities to mammalian gasdermins that mediate pyroptosis (a form of programmed necrosis) (Liu et al., 2016; Ruan et al., 2018). Multiple additional cell death mechanisms with compelling empirical evidence have been identified in bacteria and filamentous fungi, with and without parallels in animals (Erez et al., 2017; Goncalves et al., 2017; Heller et al., 2018; Smith et al., 2020). However, an equivalently advanced understanding of the dying processes in unicellular eukaryotes has not been demonstrated. Early evidence for yeast metacaspases and other components resembling mammalian apoptosis machinery were impactful by stimulating the field, but they have not garnered molecular support (Aouacheria et al., 2018; Minina et al., 2020). Thus, despite the arguments for altruistic death in unicellular eukaryotes, the question is unsettled.

Yeast are widely used models for a range of disciplines (e.g. autophagy, vesicle trafficking, cancer, neurodegeneration) and are ideal models for dissecting unicellular eukaryotic cell death. Many *Saccharomyces cerevisiae* genes have been reported to promote or inhibit death (Chaves et al., 2021), some with semblances to mammals (Fannjiang et al., 2004; Gao et al., 2019; Ivanovska and Hardwick, 2005). A common theme shared by the four best-characterized mammalian cell death pathways is membrane permeabilization carried out by pore-forming proteins such as BAX (apoptosis), MLKL (necroptosis), and gasdermins (pyroptosis), or by lipid peroxidation (ferroptosis), which typically mark a commitment or final step to cell death. Permeabilization of the yeast vacuole/lysosome has been implicated in yeast cell death, similar to death by lysosome membrane permeabilization, LMP, in mammals, but again the details are unknown (Eastwood et al., 2013; Kim and Cunningham, 2015; Kim et al., 2012; Watson and Khaled, 2020).

Surprisingly few yeast genetic screens have been aimed at identifying yeast genes that may contribute to cell death following stress (Dong et al., 2017; Jarolim et al., 2013; Kim et al., 2012; Sousa et al., 2013; Teng et al., 2011; Teng et al., 2013). Instead, more efforts have focused on yeast genes that promote drug resistance and survival, seeking to control pathogens and support wine-making industries (Todd and Selmecki, 2020; Velazquez et al., 2016). Thus, the death of drug-treated fungal pathogens are generally not studied from the perspective of understanding molecular dying processes despite public health relevance. One example of a pathogenic yeast is *Cryptococcus neoformans*, which is a major threat for HIV-infected individuals, worsened by expanding fluconazole resistance with the advent of prophylactic usage (Stott et al., 2021). Identifying mechanisms of cell death in fungi could lead to the discovery of new antifungal therapies.

Yeast may have multiple unconventional cell death mechanisms. Whether these were selected as true adaptations during evolution, or whether they can be harnessed for therapy, analogous to the anti-cancer BCL-2 inhibitor venetoclax (Roberts et al., 2016), is not yet known. Here we provide evidence for a regulated cell death pathway in *Saccharomyces cerevisiae*, and potentially in *Cryptococcus neoformans*. We propose a model where membrane-associated proteins delivered to the yeast vacuole/lysosome by AP-3 are triggered to permeabilize permeabilize the membrane and cause cell death in response to stress.

## Results

### Yeast cell death occurs over hours following stress

Non-programmed, unregulated cell death is defined by the Nomenclature Committee on Cell Death as a sudden catastrophic assault (Galluzzi et al., 2018). In this case, the cell does not contribute to its own death, and death is not preventable by any action from the cell or by any therapeutic treatment. To minimize unregulated cell death by assault, yeast cultures were treated with a near-lethal stress using a tunable heat-ramp delivered with a programmable thermocycler that heats cells gradually, 30°C to 51°C over 20 min, avoiding the assault of sudden heat shock (**Figure 1A**) (Teng et al., 2011; Teng and Hardwick, 2013).

**Figure 1.**
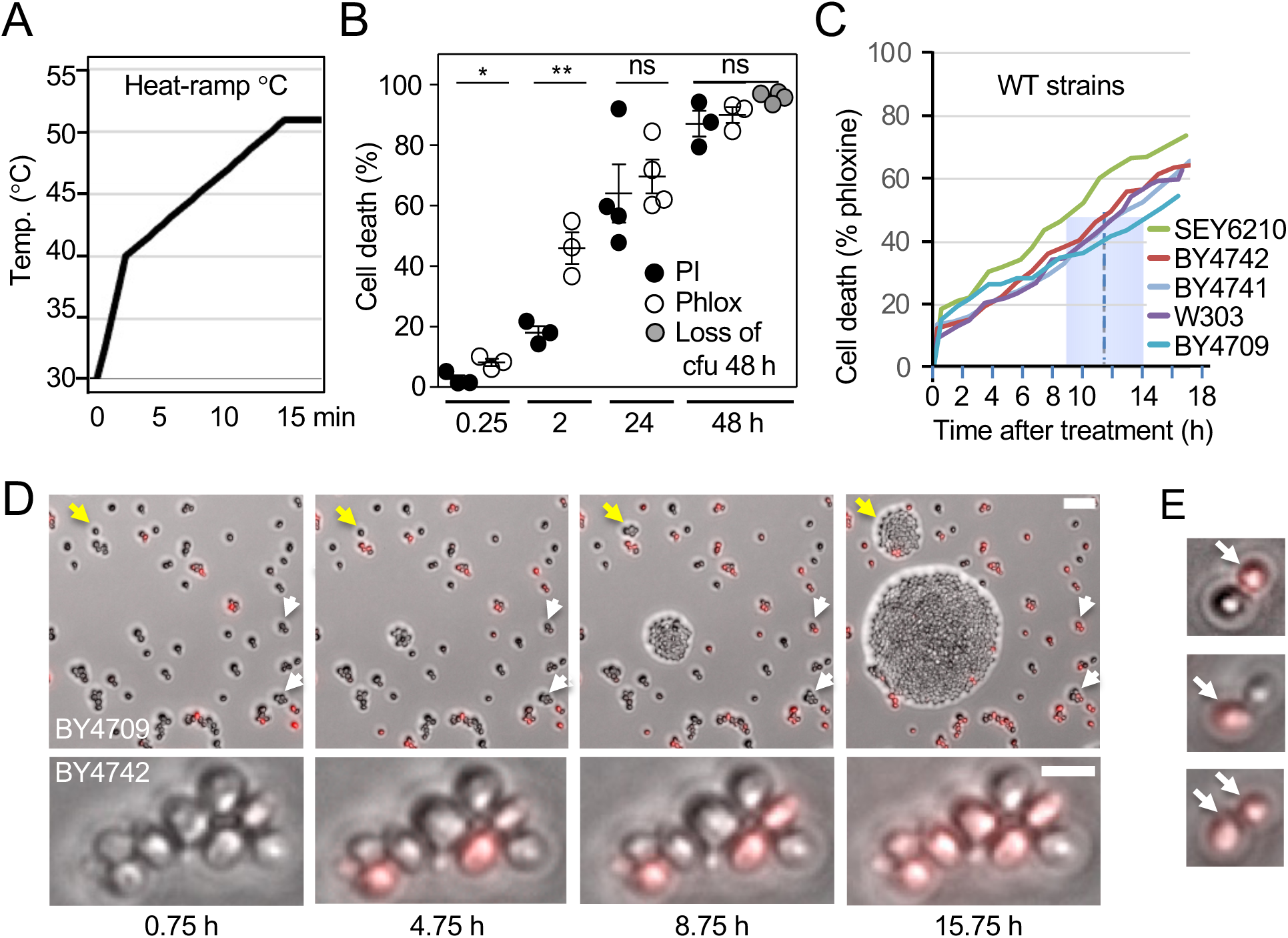
Protracted time-to-death following heat-ramp stress. (A) Temperature plot for the 20 min, 30℃-51℃ heat-ramp cell death stimulus used throughout this study to treat log phase yeast using a programmable thermocycler (Teng et al., 2011). (B) Cell death of wild type yeast BY4709 determined as percent of starting cell number stained by propidium iodide (PI) or phloxine B (Phlox) and by visible colony forming units (cfu, 48 h only, which was unaffected by the presence of dye) comparing the same samples after heat-ramp. N = 3-4 independent experiments per condition. Comparing phloxine and PI, two-tailed t-test, *p = 0.0157, **p = 0.0122, and p = 0.316 at 24 h. No significant differences between the three assay types at 48 h, two-way ANOVA with Tukey’s HSD, p = 0.1006. (C) Time-to-death determined by time-lapse microscopy of ∼300 cells per strain after heat-ramp determined by phloxine staining on YPD agar. Median time-to-death (dashed line), range (blue box) for five wild type strains. (D) Time-lapse microscopy frames for two wild type strains from an independent time-lapse experiment analyzed on agar as described in panel C. Examples of yeast cells dying at 9-16 h post heat-ramp (white arrows), and early or delayed proliferation/clonogenicity (yellow arrows). Scale bar 25 μm. (E) Examples of budded mother (lower) and daughter (upper) cell pairs stained with phloxine to detect death (arrows).

Our previous results suggested that yeast die slowly following a heat-ramp or other death stimulus (Teng et al., 2011). This conclusion was based on vital dye staining at early time points compared to clonogenic survival (colony forming units/cfu) at 2-days, with the caveat that these assays measure different cell properties. Therefore, to determine if vital dyes are useful proxies for cell death, we compared two vital dyes over an extended time course following heat-ramp, revealing that vital dyes approximate loss of clonogenic survival at 48 h (**Figure 1B**). Remaining differences are likely attributed to uncounted microscopic colonies. In addition, we observed that phloxine B, which is reportedly pumped out of living cells (Kwolek-Mirek and Zadrag-Tecza, 2014; Minois et al., 2005), detects dead/dying yeast cells several hours earlier than the DNA stain propidium iodide (PI), a field standard based on a molecule that enters cells only after plasma membrane integrity is compromised (**Figure 1B**). Using the sensitive phloxine assay, median time-to-50%-death was 11.5 h (range 9-14 h) after heat-ramp for five wild type strains (BY4709, BY4741, BY4742, W303 and SEY6210) determined by time-lapse microscopy of cells immobilized on agar (**Figure 1C**). Video frames illustrate the delay in phloxine staining (white arrows), delayed proliferation of survivors (yellow arrows), and no phloxine-positive cells reverted to phloxine-negative or began proliferating in these fields during the 16 h observation (**Figure 1D**). Death could occur first in the older mother or younger daughter cell (**Figure 1E**). Thus, yeast appear to die slowly, suggesting an ongoing molecular dying process. While this is consistent with gene-dependent cell death, it does not eliminate the possibility that cell damage incurred during heat-ramp requires many hours to manifest itself in a passive deterioration of cell function. Therefore, we took a genetic approach to identify death-promoting genes.

### Genome-wide screen identifies death-resistant AP-3 deletion strains

To identify yeast genes that promote cell death, we reanalyzed our previous genome-wide screen (Teng et al., 2011; Teng et al., 2013), this time to identify death-resistant deletion strains among ∼5000 *S. cerevisiae* knockouts (BY4741) treated with a heat-ramp (20 min, linear 30° to 62°C). Raw images of microscopic colonies in eight replicates were reacquired from a BioSpot Reader, visually inspected to remove artifacts, and systematically corrected for undercounting at increasing colony densities. A stringent cutoff yielded a hit rate of 1.84% (**Figure 2A, Supplementary Table S1**). Gene ontology function analyses of the 84 hits readily identified all four deletion strains for the small (*Δaps3*), medium (*Δapm3*), and two large subunits (*Δapl5*, *Δapl6*) of the heterotetrameric AP-3 adaptor complex involved in vesicle trafficking to the vacuole/lysosome membrane (**Figure 2B**). Our screen did not identify knockouts for AP-1 or AP-2 adaptor complexes, or the Gga1 and Gga2 adaptors that are also involved in vacuole trafficking via a separate pathway distinct from AP-3 (Buelto et al., 2020; Casler and Glick, 2020; Daboussi et al., 2012), suggesting an AP-3-specific phenotype.

**Figure 2.**
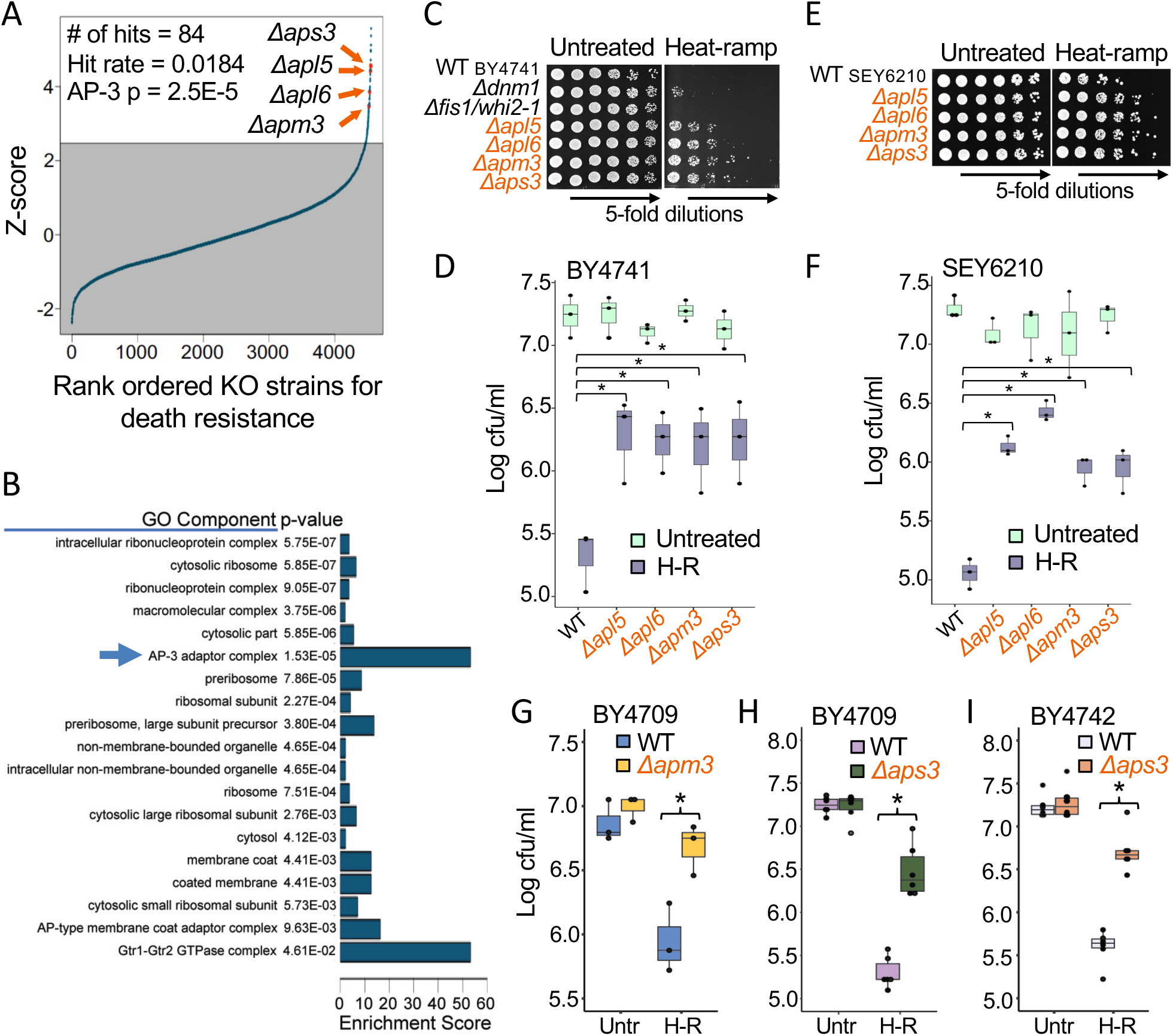
AP-3 deletion strains have greatly increased survival after heat stress. (A) Rank ordered survival of yeast knockout strains (BY4741) from genome-wide screen after heat-ramp treatment (20 min, 30°-62°C linear ramp applied to post-diauxic cells). Hit cutoff set at 1.5x interquartile range + 75% percentile of z-scores, ∼2.47. (B) Gene ontology component enrichment analysis of the 84 screen hits compared to all strains analyzed with Bonferroni correction for multiple hypothesis testing. (C) Low throughput cell death assay of log phase OD_600_-adjusted cultures (BY4741 knockout collection) spotted on plates pre and post heat-ramp. (D) Quantified data and boxplots from panel C for 3 independent experiments. Two-way ANOVA with Tukey’s HSD test post-hoc, *p ≤ 0.0048. (E and F) As described for C and D except for SEY6210 background strains. N= 3 independent experiments; two-way ANOVA with Tukey’s HSD test post-hoc, *p ≤ 0.0043. (G) Survival (cfu) of newly generated CRISPR-Cas9 deletion of *APM3* in the amino acid prototroph BY4709 pre and post heat-ramp. Three independent experiments, two-way ANOVA with Tukey’s HSD test post-hoc, *p = 0.0068. Genome sequencing and functional assays in **Supplementary Figure S1A-D**. (H) As described for panel G except for newly generated CRISPR-Cas9 deletion of *APS3* (BY4709). N= 6 independent experiments, two-way ANOVA with Tukey’s HSD test post-hoc, *p = 1.24E-08. Genome sequence and trafficking function tests in **Supplementary Figure S1E-H**. (I) As described for panel G except for new *APS3* deletion strain generated by conventional recombination in BY4742. N= 6 independent experiments, two-way ANOVA with Tukey’s HSD test post-hoc, *p = 2.4E-08.

The three heterotetrameric adaptor complexes AP-1, AP-2 and AP-3 in yeast (AP-1 to AP-5 in mammals), sort, transport, and deliver membrane-associated proteins to their respective subcellular destinations (Dell’Angelica and Bonifacino, 2019; Robinson and Bonifacino, 2001). While different AP complexes share significant sequence homology, they traffic different sets of cargo proteins in different paths with minimal overlap. Similar to mammals, yeast AP-1 and AP-3 adaptor complexes engage their respective cargo proteins by recognizing amino acid sequence motifs present in nascent, recycled or internalized proteins protruding from late/post-Golgi or endosome membranes. After engaging their cargo, AP-1 and AP-3 plus other proteins generate vesicles that transport and deliver their cargo by fusing to a target membrane with the aid of additional factors (Casler and Glick, 2020; Cowles et al., 1997a; Hirst et al., 2001; Schoppe et al., 2020; Vowels and Payne, 1998). Yeast AP-3 traffics from late/post-Golgi membranes directly to the vacuole/lysosome membrane (Cowles et al., 1997a; Odorizzi et al., 1998; Simpson et al., 1997; Stepp et al., 1997), while AP-1 is more important for Golgi and endosome recycling pathways, with a lesser role in trafficking to the vacuole (Buelto et al., 2020; Casler and Glick, 2020; Daboussi et al., 2012).

The striking death-resistant phenotype of AP-3 deletion strains (BY4741) after heat-ramp was confirmed in small scale tests (Figure 1A) and was more robust than our previous death-resistant landmark *Δdnm1* (Fannjiang et al., 2004; Ivanovska and Hardwick, 2005) (**Figure 2C and D**). Similar results were observed for the four AP-3 deletion strains in the SEY6210 background strains (**Figure 2E and F**), and for newly constructed knockouts including a CRISPR-Cas9 deletion of *APM3* (**Figure 2G, Supplementary Figure S1A-D**) and four *APS3* deletion strains engineered by different methods in two different backgrounds (**Figure 2H and I, Supplementary Figure S1E-H**). In contrast, knockouts of AP-1 subunits (Apl2, Apl4, Apm1, Apm2, Aps1), AP-2 subunits (Apl1, Apl3, Apm4, Aps2), and monomeric clathrin adaptors (Gga1 and Gga2) were all sensitive to heat-ramp similar to wild type, consistent with screen results (**Supplementary Figure S2**).

Cell death-resistance was not dependent on cell proliferation, as vital dye staining reliably distinguished AP-3-deficient yeast from wild type within 15 min after heat-ramp by phloxine staining (**Figure 3A**), and by the less sensitive PI stain (**Figure 3B**). Time-lapse microscopy revealed that AP-3-deficient yeast recovered cell proliferation faster, and linear regression analysis predicts a more protracted 50% time-to-death (20.3 h) compared to wild type (10.3 h) (**Figure 3C, 3D**, **Supplementary Movie S1**). These findings are consistent with an ongoing death process promoted by the AP-3 complex.

**Figure 3.**
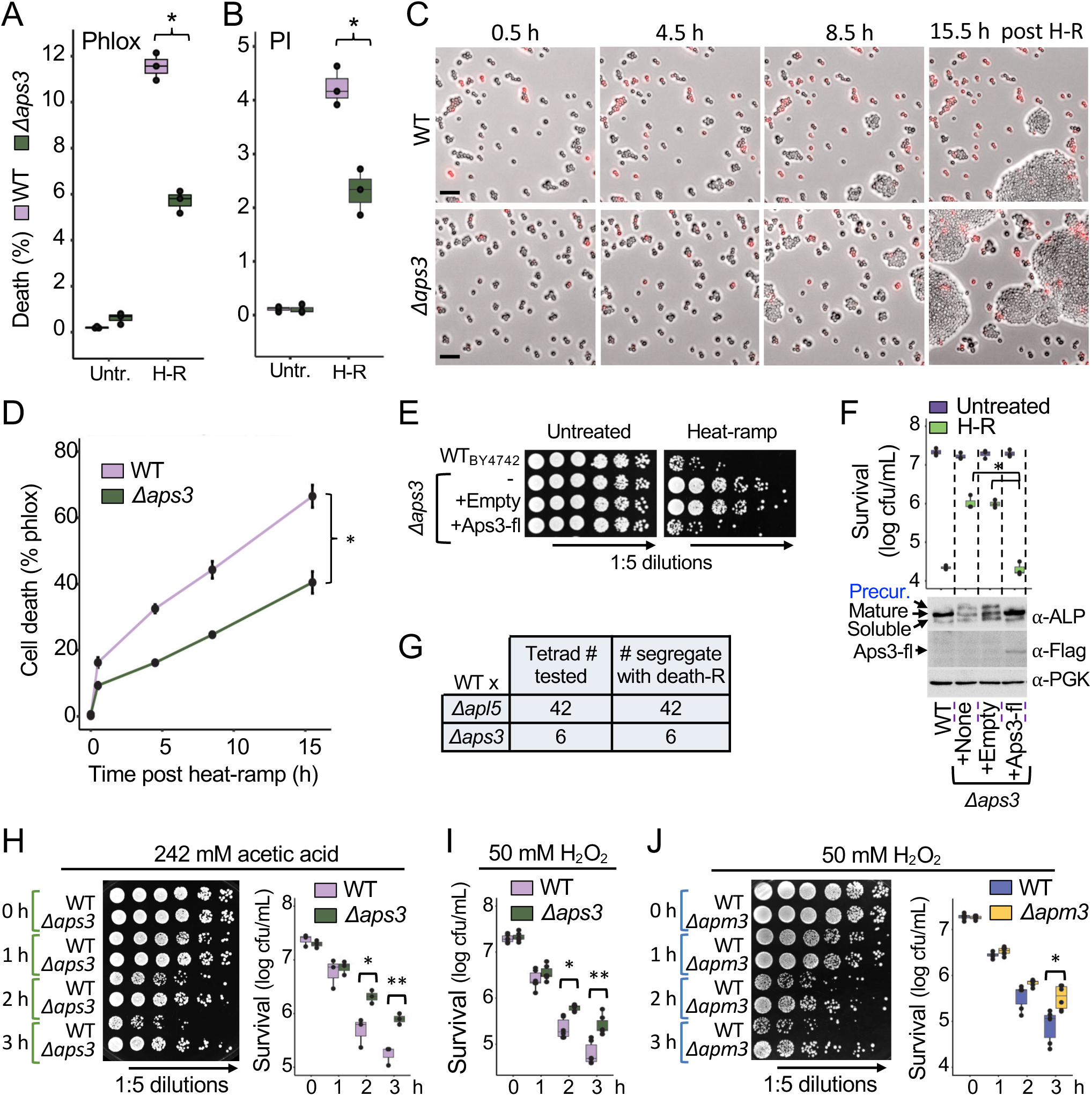
AP-3 promotes death induced by multiple stimuli. (A) Cell death of wild type and *Δaps3* determined by percent phloxine B-positive (Phlox) before and 15 min after heat-ramp using a hemocytometer. N = 3 independent experiments per condition, two-way ANOVA with Tukey’s HSD test post-hoc, *p = 4.77E-07. (B) As described for panel A except staining with propidium iodide (PI), *p = 1.54E-04. (C) Images from time-lapse microscopy showing accumulation of phloxine-positive WT and *Δ*aps3 cells after heat-ramp analyzed on YPD agar. Scale bars = 25 μm. (D) Cumulative data for 3 independent experiments described in panel C. One-way ANOVA with repeated measures, *p = 1.5E-07. (E) Rescue of *Δaps3* cell death sensitivity by expressing C-terminal flag-tagged (fl) *APS3* from its native promoter. Viability determined by cfu on selection medium before and after heat-ramp. (F) Quantified survival data for panel E. Three independent experiments, two-way ANOVA with Tukey’s HSD test post-hoc, *p = 1.53E-7 and 1.78E-07 for *Δaps3*-rescue vs. *Δaps3* +/- empty vector. WT vs. *Δaps3* (p = 1.95E-07), WT vs. *Δaps3* + empty vector (p = 2.29E-7). (G) Summary of tetrad analysis for 42 of 44 dissections yielding 42 true tetrads for all markers resulting from WT BY4742 mated to *Δapl5* and sporulated, and of all 6 analyzed tetrads for *Δaps3.* Death-resistance uniformly segregated 2:2 with the *KanMX* (KO) marker. (H) Resistance of the new CRISPR *Δaps3* strain (log phase) to 242 mM acetic acid-induced death. N=3 independent experiments, two-way ANOVA with Tukey’s HSD test post-hoc, *p = 0.0064, **p = 0.0043. (I) Resistance of *Δaps3* (CRISPR KO strain in log phase) to 50 mM H_2_O_2_-induced death. N=6 biological replicates from 3 independent experiments, two-way ANOVA with Tukey’s HSD test post-hoc, *p = 0.0014, **p = 7.95E-06. (J) Resistance of *Δapm3* (CRISPR KO strain) to 50 mM H_2_O_2_-induced death. N=6 biological replicates from 3 independent experiments, two-way ANOVA with Tukey’s HSD test post-hoc, *p = 3.25E-05.

### Cell death induced by multiple stimuli is AP-3 dependent

We previously demonstrated that 60-70% of yeast knockout strains have evolved a second gene mutation strongly affecting cell death/survival following stress, and that independent knockouts of the same gene (or genes of the same protein complex) tend to acquire mutations in a shared second gene (Cheng et al., 2008; Teng et al., 2013), analogous to *Δsod1* strains that repeatedly develop mutations in *PMR1* (Lapinskas et al., 1995). Therefore, death-resistance of AP-3 knockouts could potentially be explained by natural selection for one or more second gene mutations that arose as a consequence of deleting any one of the AP-3 genes.

However, this was not the case because cell death susceptibility was rescued by reinserting *APS3* with its native promoter into the *Δaps3* deletion strain (**Figure 3E and F**). Successful rescue of AP-3-dependent vesicle trafficking in unstressed cells was verified by monitoring the maturation of the AP-3 cargo protein alkaline phosphatase (ALP/Pho8), which defines the AP-3 pathway (also known as the ALP pathway). In the absence of AP-3, ALP/Pho8 is shunted to the vacuole by an alternate default pathway, impairing its normal maturation on the vacuole membrane by lumenal proteases that cleave the ALP/Pho8 precursor to generate mature and soluble forms (Cowles et al., 1997b; Klionsky and Emr, 1989). Using this traditional assay, we found that reinsertion of *APS3* prevented the accumulation of unprocessed ALP/Pho8 precursor (**Figure 3F**). In further support of AP-3-dependent cell death, analysis of tetrad spore sets derived from *Δapl5* and *Δaps3* backcrossed to wild type (BY4742 or other strains) resulted in 100% co-segregation of death-resistance with the AP-3 locus (*KanMX* replacing AP-3 genes) (**Figure 3G**). We conclude that AP-3, and not confounding secondary gene mutations, can promote yeast cell death following stress.

Resistance to cell death was not limited to thermal stress as disruption of AP-3 also conferred resistance to acetic acid, a biproduct of alcohol fermentation known to trigger yeast cell death, unlike other acids (Sousa et al., 2013; Vilela-Moura et al., 2011) (**Figure 3H**), and to hydrogen peroxide, a mimic of oxidative bursts produced by phagocytic host immune cells (**Figure. 3I and 3J**). Consistent with our findings, several AP-3 deletion strains were scored as death-resistant in supplementary tables of yeast cell death screens using acetic acid (Sousa et al., 2013), ER-stress (Kim et al., 2012), or thiosemicarbazone Ni(S-tcitr)2 to trigger cell death (Baruffini et al., 2020). Thus, AP-3 appears to contribute to cell death induced by multiple stimuli.

### Vesicle trafficking function of AP-3 promotes cell death

Our genome-wide screen, plus additional analyses of *Δapl5*, *Δapm3*, and *Δaps3* described thus far, imply that the AP-3 complex, rather than its individual components, can promote yeast cell death. This further implicates its vesicle trafficking function in the dying process following stress. To test this more directly, we investigated the requirement for Arf1, a small GTPase known to be required by human/yeast AP-1 and AP-3 complexes (inferred for yeast AP-3) for docking onto donor membranes to collect cargo (Anand et al., 2009; Nie et al., 2003; Ooi et al., 1998; Schoppe et al., 2020; Seaman et al., 1996). To address the role of yeast Arf1 without disrupting other membrane trafficking pathways requiring Arf1, we designed point mutations in AP-3 to prevent binding to Arf1. Mutations were selected based on sequence homology with human orthologs and a crystal structure with supporting biochemical evidence for a human-mouse hybrid AP-1 complex bound to human Arf1-GTP (PDB: 4HMY). In this structure, one Arf1 molecule is bound on the edge of each of the two large AP-1 subunits (Morris et al., 2018; Ren et al., 2013). Two amino acid changes in human AP-1β1 (corresponding to yeast Apl6 L117D/I120D in AP-3) were shown to abolish Arf1 binding and suppress Golgi localization without affecting heterotetrameric complex assembly, while mutations in the second Arf1-binding site on mouse AP-1γ1 (yeast AP-3 subunit Apl5) were only partially disruptive (Ren et al., 2013). Therefore, the Apl6 L117D/I120D changes were inserted into the yeast genome using CRISPR and sequence validated (**Figure 4A**, **Supplementary Figure S3**).

**Figure 4.**
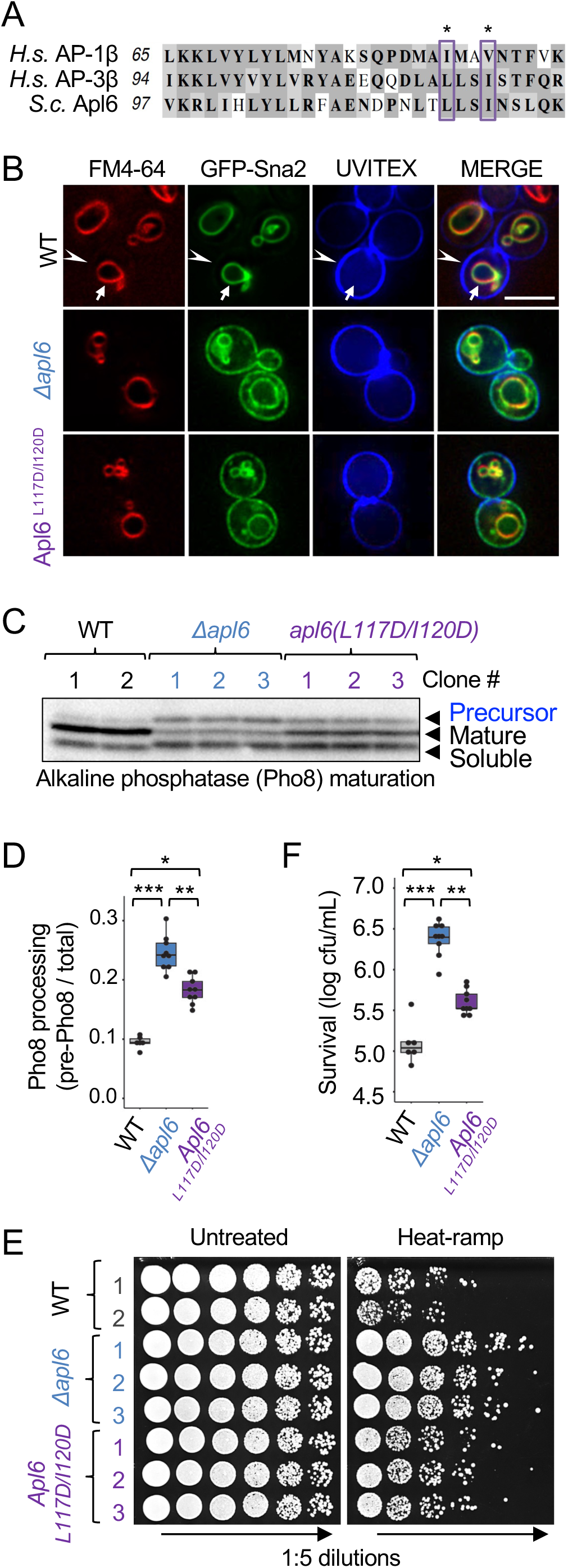
Restriction of viability relies on canonical trafficking functions. (A) Amino acid sequence alignment of yeast AP-3 subunit Alp6 with human counterparts in AP-1 (AP1β) and AP-3 (AP3β). Boxes mark the Arf1 binding sites from Ren *et al*. (Ren et al., 2013). (B) Fluorescence microscopy to monitor GFP-Sna2 localization/trafficking in the indicated yeast strains also stained with FM4-64 (red) to mark the vacuole membrane and with cell wall stain UVITEX (blue) to approximate the plasma membrane. Scale bar = 5 μm. This WT strain was transformed with the parent iCas9 and tracrRNA plasmid pCRCT (Addgene #60621) in parallel with the construction of *Δapl6* and *apl6^L117D/I120D^* for CRISPR modifications. (C) Immunoblot of endogenous alkaline phosphatase (ALP/Pho8) to detect vacuole-dependent processing of the ALP/Pho8 precursor to proteolytically matured forms. (D) Quantification for panel C, calculated as the amount of Pho8 precursor protein relative to total (precursor+mature+soluble) for 3 independent experiments after pooling results for 2-3 independent strains per genotype. Two-way ANOVA with Tukey’s HSD test post-hoc, *p = 1.56E-06, **p =3.23E-05, ***p = 1.57E-10. (E) Viability of the CRISPR modified *APL6*-modified yeast strains following heat-ramp (20 min, 30°-51°C) and spotted on agar plates. All strains were verified by Sanger sequencing (**Supplementary Figure S3**). (F) Quantified data for panel E. Viability following heat-ramp is plotted as log_10_ cfu/mL for 3 independent experiments after pooling results for 2-3 independent strains per genotype. Two-way ANOVA with Tukey’s HSD test post-hoc, *p = 2.67E-06, **p =3.98E-12, ***p= 1.06E-12.

As predicted, the Apl6(L117D/I120D) mutant strain was defective for vesicle trafficking when tested with the AP-1/AP-3 cargo reporter GFP-Sna2 (Renard et al., 2010). Normally present on the vacuole membrane (marked with FM4-64) in wild type cells, GFP-Sna2 was mislocalized to the plasma membrane (approximated by cell wall stain UVITEX) in the *apl6^L117D/I120D^* mutant, similar to the knockout *Δapl6* (**Figure 4B**). These results were confirmed for the endogenous AP-3 cargo protein ALP/Pho8 based on accumulation of its precursor form in *apl6^L117D/I120D^*, although to a lesser extent compared to *Δapl6*, indicating partially defective AP-3 trafficking in the mutant *apl6^L117D/I120D^* (**Figure 4C and D**). Importantly, the pro-death function of AP-3 was diminished by ∼85% by the Arf1 binding site mutations on Apl6 (*apl6^L117D/I120D^*) in three independent strains following heat-ramp (**Figure 4E and F**). Thus, specific disruption of the trafficking function of AP-3 also impairs cell death. A second Arf1 binding site on Apl5 could potentially account for the incomplete effects of Apl6(L117D/I120D) (Morris et al., 2018; Ren et al., 2013). These findings indicate that a conserved Arf1 binding site on yeast Apl6 is required both for normal vesicle trafficking and for stress-induced yeast cell death.

### AP-3 is required shortly before the death stimulus

The AP-3 complex could potentially exhibit a gain of pro-death function to act as an activated direct effector of cell death under stress conditions. However, this model is challenged by the unlikely possibility that the large 275,000 kDa AP-3 tetramer is thermostable under heat-ramp conditions (Leuenberger et al., 2017). We also disfavor the possibility that heat-induced malfunction of AP-3, or the resulting mislocalization of its cargo, promotes cell death, as these scenarios are more in line with the AP-3 knockout condition (which survive better). Instead, we favor a model where AP-3 is the delivery vehicle for cargo proteins that exhibit stress-induced toxic effects resulting in vacuole membrane damage. In this model, AP-3 is predicted to deliver its death-promoting payload to the vacuole prior to a death stimulus.

To investigate this possibility, the auxin-induced degron (AID) system (Morawska and Ulrich, 2013; Nishimura and Kanemaki, 2014) was used to inactivate AP-3 rapidly. Strains were engineered with an AID-6xFlag cassette fused to the C-terminal knockin TAP-tag to generate endogenous AID-tagged AP-3 subunits (Snyder et al., 2019). This cassette also encodes the auxin-responsive OsTIR1 cullin adaptor needed to recruit AID-tagged proteins to the yeast cullin-1 E3 ligase for ubiquitination and degradation. Of the four AID-tagged AP-3_TAP_ strains, Apl5 (Apl5_TAP_-AID-flag) was the most stably expressed subunit and was used for testing thereafter. Importantly, only the AID-tagged strain was protected from heat-ramp-induced death when pre-treated 4 h with auxin (50 μM IAA) to degrade Apl5, restoring survival to a level similar to the *apl5* knockout (**Figure 5A and 5B**). Reduced baseline survival after heat-ramp of the Apl5_TAP_-AID strain relative to the Apl5_TAP_ control is likely due to leaky auxin-independent activities of OsTIR as reported (Yesbolatova et al., 2020). Together these findings indicate that AP-3 is required prior to the death stimulus.

**Figure 5.**
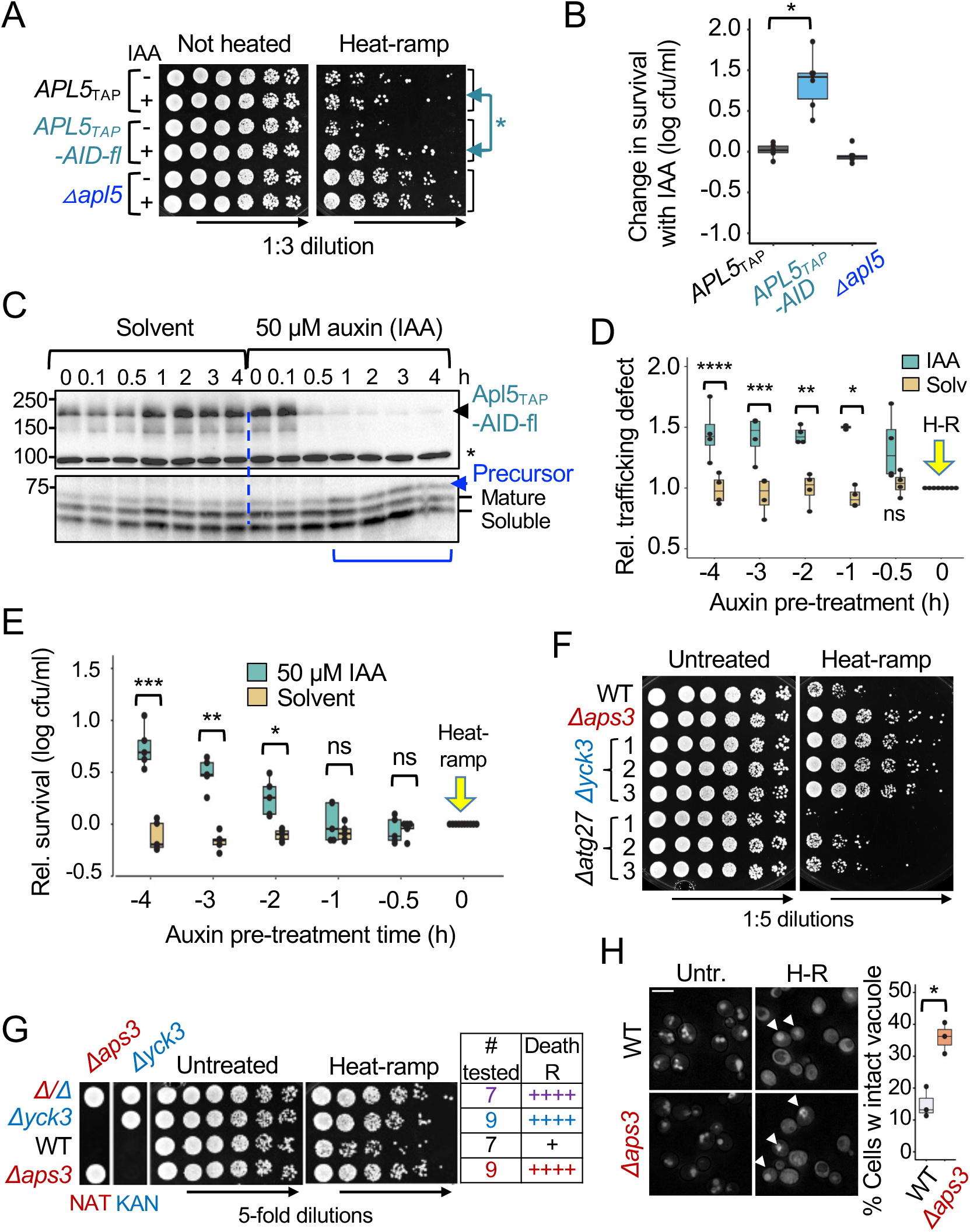
AP-3 is required shortly before the cell death stimulus. (A) Effect of transient degradation of AP-3 determined by survival of yeast preincubated 4 h with either solvent control (ethanol) or 50 μM auxin (IAA) to degrade Apl5-TAP-AID-6xflag, and plated in 3-fold serial dilutions before and after heat-ramp (20 min, 30°-51°C). (B) Quantified data for panel A for 6 independent experiments plotted as [(log_10_ cfu/mL with auxin) - (log_10_ cfu/mL with ethanol)]. Two-way ANOVA with Tukey’s HSD test post-hoc, *p= 6.81E-07. Comparing *Δapl5* to either *APL5_TAP_-AID* (p = 0.931) or to *APL5_TAP_-AID* (p = 2.2E-07). (C) Immunoblots of the *APL5-AID* strain to monitor the degradation of Apl5-TAP-AID-6xflag during 0-4 h incubation with either solvent control (ethanol) or 50 μM auxin (IAA) detected with anti-flag (upper), and AP-3 function was monitored by maturation status of Pho8 detected with anti-alkaline phosphatase (lower). *Non-specific band. (D) Quantification for panel C. ALP/Pho8 defect was calculated from densitometry as amount of precursor protein relative to total (precursor+mature+soluble), and relative ALP/Pho8 defect was plotted as [(ALP defect with auxin or ethanol)/(ALP defect of untreated)] for 4 independent experiments. Two-way ANOVA with Tukey’s HSD test post-hoc, ****p = 9.5E-4, ***p = 5.7E-4, **p = 0.00201, *p = 2.09E-5, ns = 0.128. (E) Enhanced yeast survival requires degradation of Apl5-AID (Apl5-TAP-AID-6xflag) 1-2 h prior to a death stimulus. Relative survival calculation [(log_10_ cfu/mL with auxin or ethanol) - (log_10_ cfu/mL untreated)] for samples pre-incubated with either solvent control (ethanol) or 50 μM auxin (IAA) prior to heat-ramp (20 min, 30°-51°C) at time = 0 (yellow arrow). Auxin/ethanol-containing samples were plated and colonies enumerated at 48 h. N = 5 independent experiments, two-way ANOVA with Tukey’s HSD test post-hoc, *p = 0.0016, **p = 5.2E-9, ***p = <1.0E-13. (F) Heat-ramp cell death assay (20 min, 30°-51°C) for three single cell-derived substrains (BY4741 KO collection) corresponding to AP-3 cargo proteins Yck3 and Atg27. (G) Survival of plated yeast after heat-ramp (20 min, 30°-51°C), and NAT or KAN (G418) drug selectivity are shown for a representative tetrad resulting from crossing *Δaps3*::*NatMx6* with *Δyck3*::*KanMx4.* The number of spore-derived strains tested (total of 8 tetrads) and their heat-ramp phenotypes are indicated. (H) Wild type and *Δaps3* cells were pre-stained with vacuole dye CMAC, and unstressed and heat-ramp-treated samples were imaged in rich (YPD) medium at 30°C. Two-tailed t-test for 3 fields from one experiment, counting >1500 cells per genotype, *p = 0.006319.

To determine how long AP-3 is needed prior to the death stimulus, a time course of progressively shorter auxin pre-incubation times were compared. Apl5-AID protein was maximally degraded by ∼30 min after addition of auxin (50 μM IAA) (**Figure 5C** **upper**), and AP-3 trafficking function was noticeably impaired by ∼1 h after auxin treatment assessed by the accumulation of unprocessed ALP/Pho8 precursor (**Figure 5C and 5D**). For cell death, AP-3 was required for at least 1-2 h prior to heat-ramp to stimulate significant cell death, as degradation of Apl5 within the hour prior to the death stimulus did not significantly impair cell death (**Figure 5E**). This supports the model that AP-3 delivers sufficient amounts of its latent deadly cargo to the vacuole by 2 h before the death stimulus to initiate cell death. Thus, AP-3 is a critical component of this death pathway, but is not necessarily a direct effector of death.

### AP-3 cargo protein Yck3 is a death effector of AP-3 at the vacuole

AP-3 cargo proteins are prime candidate mediators of cell death. AP-3 is likely a substantial contributor to the vacuole membrane proteome, yet relatively few AP-3 cargo proteins have been identified, and none of these are among the 84 hits from our screen. However, because most yeast knockouts contain a secondary mutation in at least 20% of their cell population that strongly affects cell death, typically increasing death (Teng et al., 2013), we tested single cell-derived colonies/substrains from each knockout for ten confirmed AP-3 cargo proteins: ALP/Pho8, the protein kinase Yck3 (Sun et al., 2004), a vacuole tSNARE Vam3 (Cowles et al., 1997a) and vSNARE Nyv1 (Darsow et al., 1998), amino acid transporters Ypq1-3 (Llinares et al., 2015), a transmembrane component of ESCRT-0 also involved in phagophore formation Atg27 (Segarra et al., 2015), the AP-3 reporter proteins Sna2 (Renard et al., 2010) and Sna4 (Pokrzywa et al., 2009) of unclarified functions, and two candidate AP-3 cargo Cot1 and Zrt3 (Yang et al., 2020). Small scale validation experiments confirmed the genome-wide screen results for all candidates tested with the exception of *Δyck3* substrains, which were strikingly death-resistant similar to AP-3 knockouts in the log phase heat-ramp assay from figure 1A (**Figure 5F, Supplementary Figure S1 and S4**). In contrast, the same *Δyck3* substrains were death-sensitive in the post-diauxic growth phase heat-ramp assay used for the genome-wide screen (**Supplementary Figure S5A**). In addition, this discrepancy is due to a secondary gene mutation present in *Δyck3* affecting survival only in the post-diauxic assay based on backcrossing to WT (and *Δaps3*) and tetrad analysis (**Figure 5G, Supplementary Figure S5B and S5C**). Thus, loss of *YCK3* appears to be responsible for death-resistance, indicating pro-death effects of Yck3 following stress.

Consistent with the AP-3 complex and the Yck3 kinase working together in the same trafficking pathway, they also appear to work together in the same cell death pathway following stress, as double knockouts (*Δaps3/Δyck3*) phenocopy single knockouts (**Figure 5G, Supplementary Figure S5C**). Interestingly, endogenous and expressed Yck3 kinase was reported by others to promote loss of yeast viability after heat stress (Karim et al., 2018).

### Vacuole membrane permeabilization occurs long before early markers of cell death

Vacuole/lysosome membrane permeabilization has been suggested to be a key event during yeast cell death (Chaves et al., 2021; Eastwood et al., 2012; Eastwood and Meneghini, 2015; Kim and Cunningham, 2015; Kim et al., 2012). The yeast vacuole is a large organelle that can make up a quarter of the cell volume (Klionsky and Eskelinen, 2014). Thus, it is expected that consequential disruption of pH and redox states and the release of degradative enzymes or other events upon leaking vacuolar contents could lead to cell demise. However, detailed mechanisms of vacuole membrane permeabilization and whether this event is a facilitator or consequence of the dying process is more challenging to address. To determine if AP-3 promotes loss of vacuole membrane integrity following stress, cells were stained prior to heat-ramp with the thiol-reactive dye CMAC-blue (210 MolWt). CMAC accumulates in vacuoles and fluoresces in response to the reducing environment, but can leak into the cytoplasm if the vacuolar membrane is compromised (Karim et al., 2018; Llinares et al., 2015). In unstressed cells, CMAC was uniformly restricted to vacuoles (**Figure 5H**). However, vacuoles of both wild type and *Δaps3* likely underwent homotypic fusion during heat-ramp as cells tended to have a single large vacuole, consistent with previous reports of protective vacuole stress responses after heat (Schoppe et al., 2020). In support of our model, fewer wild type cells retained vacuolar CMAC compared to *Δaps3* at 30 min post heat-ramp (**Figure 5H**). Thus, AP-3 can promote permeabilization of the vacuole membrane following stress.

To further address the role of vacuole membrane permeabilization, we determined if permeabilization is an early or late event in the dying process by monitoring the timing of CMAC leaking into the cytoplasm relative to cell death following heat-ramp. CMAC release was consistently observed in cells before the first hints of phloxine staining, an early marker of cell death. In the example cell, CMAC begins to leak at ∼6.0 h after heat-ramp (**Figure 6A**, arrows in heatmap) and released CMAC gradually diffused throughout the cell by 7.0 h (green fluorescence) (**Figure 6A** **and Supplementary Movie S2**). Not until approximately 3 h after CMAC began leaking from the vacuole, the early death marker phloxine became detectable in the cytoplasm (**Figure 6A**, arrowheads), which gradually accumulated over the next 30 min. From figure 1B, the plasma membrane is not expected to become permeable to propidium iodide for several more hours after phloxine staining. Cells that die at earlier times after stress can also have similar delays between CMAC leakage and phloxine staining (**Supplementary Figureure S6A**), although this timing can vary considerably, similar to mammalian cell death following treatment with the death agonist TRAIL (Spencer et al., 2009).

**Figure 6.**
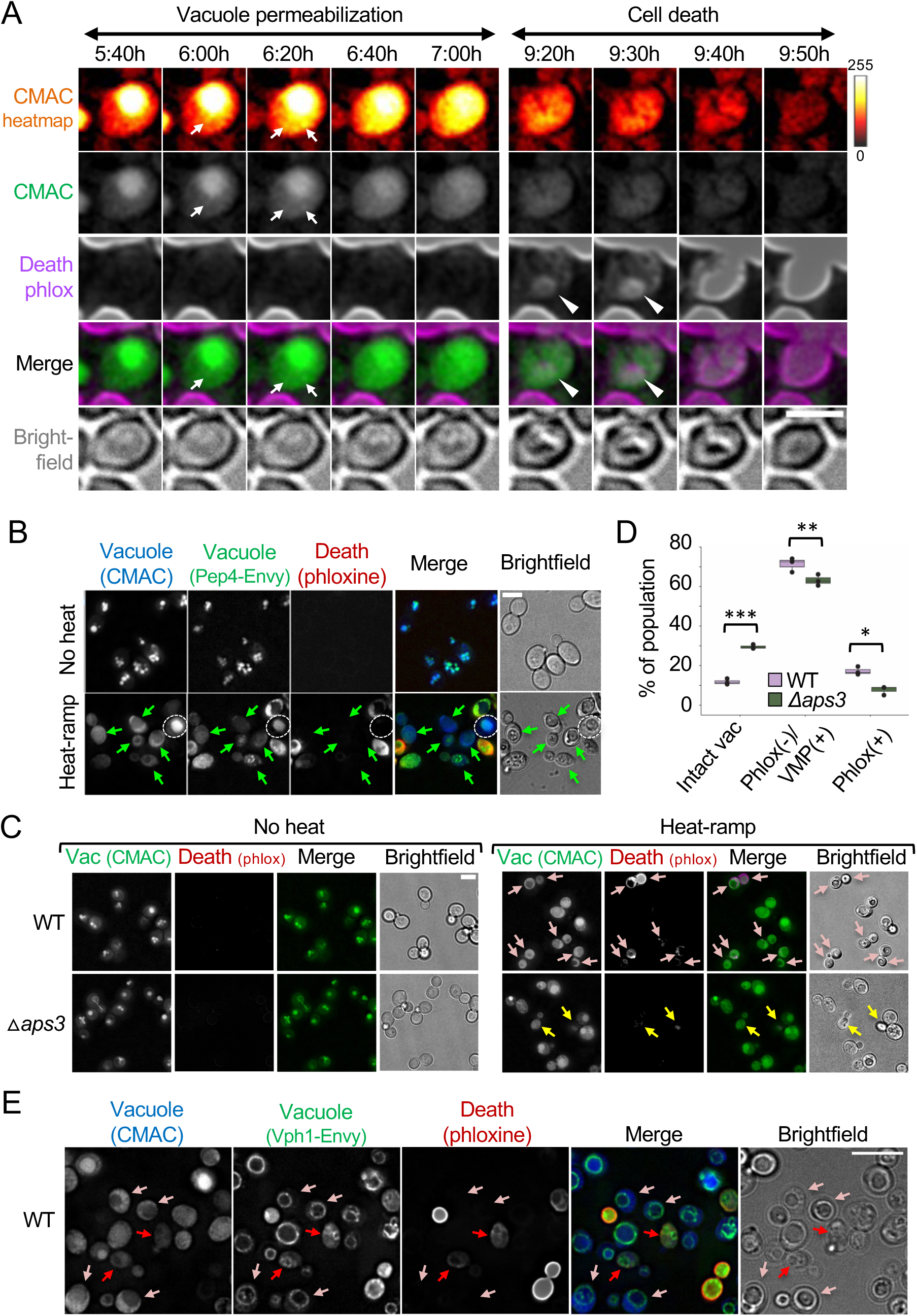
AP-3-dependent vacuole membrane permeabilization. (A) Time-lapse microscopy of wild type yeast cells stained with CMAC prior to heat-ramp (20 min, 30°-51°C), and imaged every 10 min in rich (YPD) medium containing 2 μg/mL phloxine at 30°C. CMAC is presented as pixel intensity heatmap (color key on right), gray scale and green fluorescence. Scale bar 2.5 μm. (B) Yeast expressing genomic *PEP4-ENVY* were pre-stained with CMAC and heat-ramp-treated as in panel A, except cells were transitioned to synthetic complete medium (SC^AUX^, to avoid autofluorescence) (Teng et al., 2018) containing 2 μg/mL phloxine at 30°C immediately before imaging at ∼20 min after heat-ramp. Arrows mark examples of live (phloxine-negative) cells with vacuole markers CMAC and Pep4-Envy present in the cytoplasm. Scale bar 5 μm. (C) WT and *Δaps3* cells were pre-stained with CMAC prior to heat ramp and imaged in YPD medium containing 2 μg/mL phloxine at 20-30 minutes after heat-ramp treatment. Representative images presented. Scale bar 5 μm. (D) Quantification for panel C. Each cell imaged was classified as phloxine-positive (Phlox+), phloxine-negative with permeabilized vacuole membranes (Phlox-/VMP+), or intact vacuoles (Vac). Two-way ANOVA with Tukey’s HSD test post-hoc for 3 fields from one experiment, counting >300 cells per genotype, *p = 0.0035, **p = 0.012, ***p = 1.05E-5. (E) Yeast cells expressing genomic *VPH1-ENVY* were imaged as in panel B. Rose arrows mark dying cells that have cytoplasmic CMAC staining but retain vacuole organelle integrity (VPH1-ENVY) but no detectable phloxine dye staining. Red arrows mark cells in the process of staining with phloxine-positive that have undergone vacuolar membrane permeabilization. Scale bar 5 μm.

To determine if vacuole membranes also become permeable to larger molecules, we monitored localization of the endogenous vacuolar protease Pep4 with a C-terminal GFP-Envy tag (Pep4-Envy, 71.5 kDa). We observed non-vacuolar cytoplasmic Pep4-Envy in live phloxine-negative cells after heat-ramp, and only in cells with CMAC staining outside the vacuole (**Figure 6B**, arrows). Thus, the pores that develop in the vacuole membrane are sufficiently large to pass intact proteins. These results indicate that stress-induced release of vacuolar contents into the cytoplasm occurs before early detection of impending cell death, and further suggests that this is a key event leading to cell death.

### AP-3-dependent vacuole membrane permeabilization during cell death

To evaluate the effects of AP-3 on vacuole membrane permeabilization relative to cell death at the single cell level, cells were prestained with the vacuolar dye CMAC before heat-ramp and cell death was assessed with phloxine. We also found that dead/dying (phloxine-positive) wild type cells typically have stronger cytoplasmic-to-vacuole CMAC signals, indicative of compromised vacuole membrane integrity in dying cells (**rose arrows, 6C**). In contrast, phloxine-positive *Δaps3* cells have an inverse ratio of cytoplasmic-to-vacuole CMAC (**yellow arrows, 6C**), supporting the model that AP-3 promotes vacuole membrane permeabilization during cell death. Importantly, of the cells that had leaked CMAC into the cytoplasm, fewer *Δaps3* cells were also stained with phloxine compared to wild type (**Figure 6D**). However, vacuole permeability was not completely blocked in AP-3-deficient cells, potentially reflecting default AP-3-independent transport of pro-death cargo proteins to the vacuole, or due to alternative molecular dying processes.

To further assess the structural integrity of vacuole organelles following membrane permeabilization, vacuole morphology was monitored in yeast expressing GFPEnvy fused to the C-terminus of endogenous Vph1, a subunit of the vacuole membrane-embedded Vo domain of the V_1_Vo-ATPase. Vph1-Envy rings marking the vacuole membrane remain clearly visible in cells that have fully or partially released CMAC from the vacuole (**rose arrows**, **6E**), and the vacuolar membrane remains visible even in cells at the early/dim stages of phloxine staining (**red arrows**, **6E**). Conversely, the vacuole membrane is no longer detectable in cells brightly stained with phloxine. Thus, vacuole membranes become leaky before loss of organelle integrity. Localized membrane disruptions are consistent with microscopy imaging in figure 6A (arrows). The mechanism of permeabilization could be non-specific (e.g. lack of membrane repair, unfolded integral membrane proteins). However, the general trend in the mammalian cell death field is that death-promoting, pore-forming molecules continue to be identified, suggesting the specificity of these processes. For example, 16 kDa human/mouse NINJ1 oligomerizes and ruptures the large plasma membrane balloons that emerge on dying mammalian cells in culture, previously assumed to be a passive event (Kayagaki et al., 2021). In contrast, Ninj1 is required to release proinflammatory DAMPs (but not IL-1β) during pyroptosis (Kayagaki et al., 2021). Our results in yeast are consistent with selective vacuole membrane permeabilization as a key step in the dying process.

### Potential conservation of an AP-3-dependent cell death pathway

To test the possibility that AP-3 may promote cell death in other yeast species, we obtained the deletion strain of CNAG_02468 encoding *Cryptococcus neoformans* Ap3d1, homolog of *Saccharomyces cerevisiae* Apl5. Using a modified heat-ramp assay to accommodate the known heat-sensitivity of *Cryptococcus* species, we found that *C.n. Δapl5*, was strikingly death-resistant compared to wild type control (**Figure 7A and B**).

**Figure 7.**
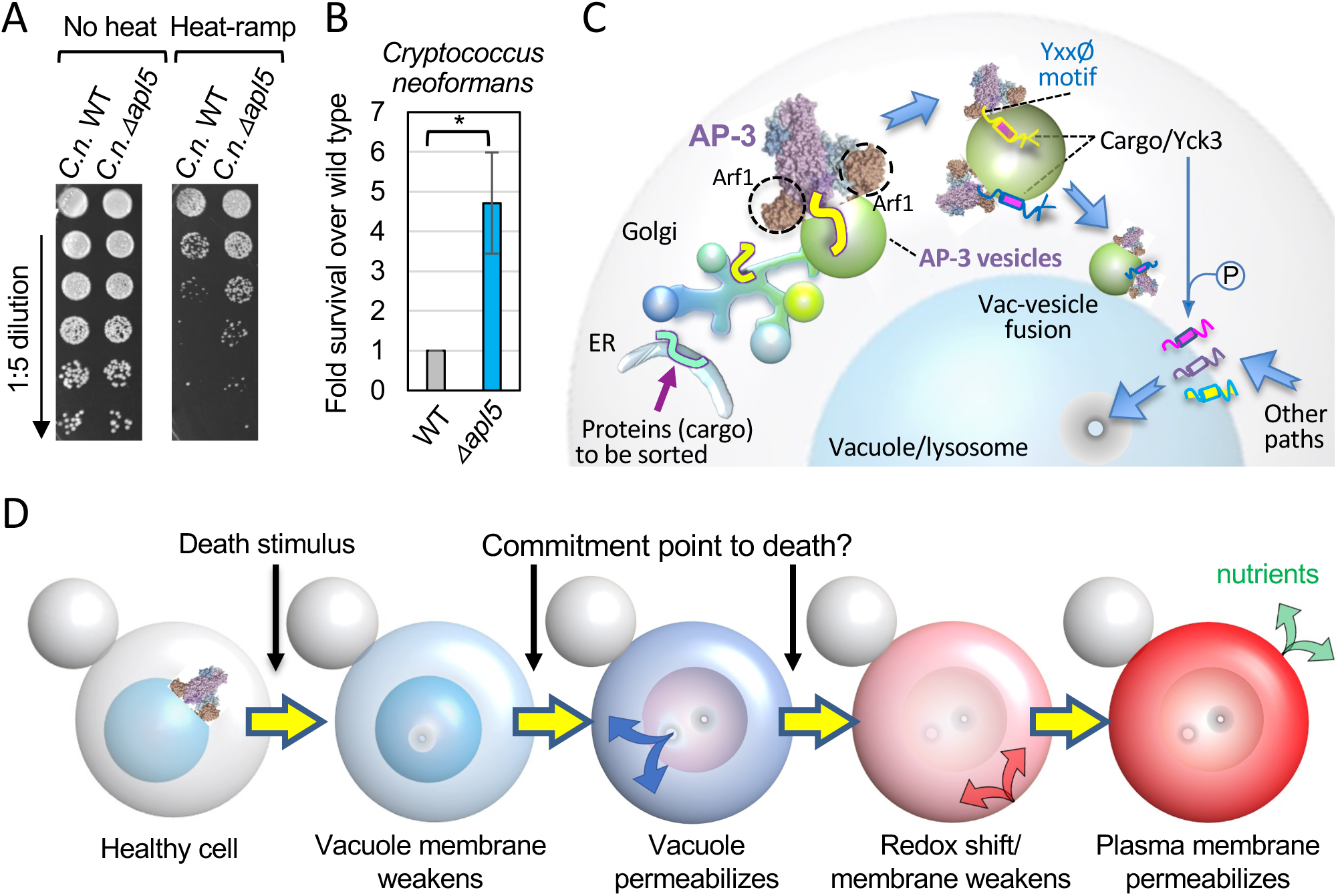
*Cryptococcus neoformans* AP-3-deficiency enhances survival. **(A)** Heat-ramp cell death assay adapted for *Cryptococcus neoformans*, comparing wild type and AP-3 knockout of the Apl5 homolog (CNAG_02468) for survival. (B) Quantified data for A, n=4 biological replicates on the knockout population and two subcolonies tested in 2 independent experiments, normalized to wild type control and presented as mean +/-SD. Two-tailed t-test, *p = 0.0177. (C) Proposed model for healthy and pro-death AP-3 vesicle trafficking pathway. Arf1-GFPase-dependent docking of AP-3 at late Golgi/trans-Golgi network (TGN) membranes (not stacked in yeast, e.g. Movie S3 of (Levi et al., 2010)) AP-3-specific recognition of newly synthesized di-leucine and tyrosine motif (Yxxϕ)-containing proteins (cargo), vesicle formation and transport, vesicle fusion to the vacuole to deliver resident vacuole membrane proteins that carry out normal cell functions, including the membrane-associated casein kinase Yck3, which phosphorylates several other membrane-associated proteins to regulate vesicle fusion, homotypic vacuole fusion, and other functions. Following cell death-inducing stresses, Yck3 and potentially other AP-3 cargo proteins and/or lipids result in permeabilization of the vacuole membrane by an unknown mechanisms, resulting in yeast cell death. (D) Sequence of events during yeast cell death in response to lethal stress. Yeast vacuoles undergo Yck3-regulated homotypic fusion (when membrane damage could potentially occur). Minutes to hours after stress, vacuolar membranes begin to leak and become permeable to CMAC blue, and several minutes-hours later the cytoplasm undergoes a gradual increase in staining by redox-sensitive phloxine B. Minutes-hours later the plasma membrane becomes permeable to propidium iodide, and the release of cellular content from dying cells may serve as a nutrient source for survivors.

Heat-resistance of *C.n.* at body temperature versus temperatures below 30 °C has been associated with virulence (Robert and Casadevall, 2009). An analysis of thermal tolerance of fungal species revealed that the majority grew well at temperatures below 30 °C but there was a rapid decrease with higher temperatures such that only a minority could survive at mammalian temperatures (Robert and Casadevall, 2009). Consequently, mammalian body temperatures create a thermal restriction zone for many potential fungal pathogens and thus provide considerable protection against mycoses. Thus, we also uncovered a new mechanism for thermal killing of fungi.

### How do yeast die in the AP-3 pathway?

We suggest a model for a molecular cell death pathway conserved in distantly related yeast species. In this model, cell death is mediated by the delivery of cargo-laden AP-3 vesicles to the vacuole membrane, where specific cargo proteins including Yck3 kinase facilitate vacuole membrane damage following stress (**Figure 7C and D**). The pro-death effects of Yck3 and presumably other factors are dependent on cell stress, but the mechanisms that activate or otherwise alter Yck3 function resulting in localized permeabilization of vacuole membranes are unknown. We suppose that the protracted time-to-death after heat-ramp is required for potentiation of pro-death effector function. AP-3-dependent release of vacuole proteases presumably digests cellular substrates in dying cells, which are predicted to facilitate the utilization of this digested material by surviving cells (**Figure 7C and D**). Analogous to mammalian cell death pathways leading to membrane disruption, vacuole membrane permeabilization in yeast may serve as the point of no return beyond which few cells survive.

## Discussion

We describe a new mechanism for thermal killing in fungi, suggesting a potential new avenue for drug discovery as drug-mediated stimulation of these cell death pathways could represent a novel strategy for antifungal drug discovery (Kulkarni et al., 2019; Robert and Casadevall, 2009).

### How does Yck3 promote cell death?

Because AP-3 and its cargo are found on both the Golgi/endosome membranes where vesicles are formed (where Arf1 is required), and also on the destination membrane where cargo are delivered (e.g. Sna2), it is challenging to distinguish which of these sites is the nexus for cell death. However, an enforced-retargeting strategy recently confirmed that Yck3 is dispensable for vesicle formation at origin membranes, and instead functions at the vacuole membrane where Yck3 promotes tethering and SNARE-mediated fusion of vesicles with vacuole membranes (Cabrera et al., 2010; Schoppe et al., 2020). These findings support our model that Yck3 kinase promotes cell death by acting at the vacuole membrane. However, the targets of Yck3 kinase that are responsible for cell death are unclear, and whether this involves a pore-forming factor that permeabilizes the vacuolar membrane is unknown.

However, AP-3 and Yck3 may facilitate cell death by an alternative mechanisms. Consistent with our findings, others found that an overactive Yck3 kinase (D44N), or Yck3 overexpression, reduces yeast viability following stress induced by cycloheximide or mild heat (Karim et al., 2018). However, they propose an alternative but not mutually exclusive mechanism of cell death. These pro-death effects of Yck3 are suggested to occur through its role in vacuole-vacuole organelle fusion. In this model, overzealous Yck3 impairs vacuole membrane turnover, thereby depriving cells of a protein quality control mechanism, and simultaneously limiting the supply of vacuole membranes and proteins as nutrient sources (Karim et al., 2018). This is because homotypic vacuole fusion is one type of protein quality control mechanism for turning over vacuole membranes and proteins that become entrapped within a membrane disc that forms were two vesicles become pressed together (McNally et al., 2017). Cycles of Yck3 kinase activity are suggested to fine-tune the hemifusion pore that forms between two vesicles. Their model proposes that active Yck3 speeds up homotypic vacuole fusion too fast to permit the formation of membrane discs, known as intralumenal fragments (ILF), carrying proteins for degradation. Conversely, more limited Yck3 activity allows expansion of the hemifusion pore that forms between two vacuoles prior to fusion, facilitating full fusion to occur along the rim of the disc, thereby releasing the membrane disc into the vacuole lumen for recycling (Karim et al., 2018). The key target of Yck3 kinase activity for these events is Vps41 in the HOPS complex. Dephosphorylated Vps41 stabilizes primed SNAREpins to delay fusion. In this scenario, Yck3 could promote cell death by suppressing turnover of misfolded proteins and damaged membranes. Other proteins phosphorylated by Yck3, such as Env7 (vacuole membrane kinase) (Manandhar et al., 2020), Mon1-Ccz1 (GTP exchange factor/GEF for Ypt7/Rab7) (Lawrence et al., 2014), and the SNARE Vam3 (Brett et al., 2008) are also potential death effectors.

Another more passive mechanism of cell death resistance by AP-3 and Yck3 knockouts is that the abundance of proteins in the vacuole membrane may be reduced compared to wild type, and in turn reduces the problem of stress-induced unfolded vacuole membrane proteins that destabilize the membrane. Thus, the need for this quality control mechanism could be reduced in the absence of AP-3. A related mechanism of mammalian cell death has been suggested, where the heavily membrane-laden mitochondrial inner membrane may undergo permeability transition when cell stress leads to misfolded polytopic proteins resulting in membrane disruption (Lemasters et al., 2009).

Yck3 kinase was previously shown to be an AP-3-specific cargo protein (Sun et al., 2004). Mutation of a consensus tyrosine motif Yxxϕ (Yck3 residue 444-YDSI) required for its interaction with AP-3 impairs its vacuole membrane localization, although like ALP/Pho8, Yck3 can reach the vacuole by default pathways (Sun et al., 2004). The involvement of Yck3 in cell death further supports the role of AP-3-dependent vesicle trafficking in cell death. Alternative paths to the vacuole may contribute to vacuole membrane protein in AP-3-deficient cells and *YCK3* knockouts.

One potential caveat to our model is that Yck3, in addition to being an AP-3 cargo protein, is also critical for the fusion of AP-3 vesicles to the vacuole by phosphorylating Vps41 in the HOPS tethering complex that regulates fusogenic SNAREs such as the AP-3 cargo proteins Vam3 and Nvy1 (Brett et al., 2008; Cabrera et al., 2009; LaGrassa and Ungermann, 2005). However, Yck3 as a cargo protein versus as a regulator of subsequent membrane fusion are separable functions. For one example, a short segment of Yck3 containing its Tyr motif (residues 409-462) is sufficient to confer AP-3-dependent trafficking to a heterologous protein (Sun et al., 2004).

## Methods

### Yeast strain generation, culturing and plasmids

Yeast strains for this study are listed in **Supplementary Table S2**, and plasmids, primers, gene blocks in **Supplementary Table S3**. Frozen *S.c.* yeast stocks were grown on YPD (2% peptone, 1% yeast extract, and 2% dextrose) agar plates for 2 days at 30°C. GFP-*SNA2* plasmids with native promoter (pRS416) were provided by Pierre Morsomme (Renard et al., 2010). Yeast were transformed as described (Gietz and Schiestl, 2007), selected and maintained on synthetic complete (SC) medium (Teng et al., 2018) minus uracil. Tetrad analysis was performed as reported (Teng et al., 2013).

Genetic rescue of *APS3* deletion strains was engineered by integrating the *APS3* gene with 133 bp 5’ UTR (distance to next gene) and C-terminal flag-tag (pRS303) into the *his3* locus. After selection in minus-histidine, strains were cultured in YPD. Envy-tagged strains were engineered by capturing GFP-Envy from Addgene plasmid #60782 using PEP4 or VPH1-specific primers as described (Slubowski et al., 2015).

CRISPR-modified strains were engineered using gBlocks (Integrated DNA Technologies/IDT) with ≥45 bp flanking the Cas9 recognition site, and inserted into the uracil pCRCT plasmid (Addgene #60621) as described (Bao et al., 2015). Transformants were maintained under selection (-uracil) for 6-days, and confirmed by genome sequencing. Synthetic complete minimal medium SC^AUX^ (0.67% yeast nitrogen base w/o amino acids, amino acids for BY4741/BY4742 auxotrophies, 2% dextrose, 20 mg/L uracil) (Teng et al., 2018) was used for tetrad analyses of CRISPR-derived strains (Teng et al., 2013). For *APL6* knockin mutations, strains were screened for loss of pCRCT plasmids prior to transformation with the cargo tracking plasmids pGFP-*SNA2* (-URA). Auxin-inducible degron (AID)-tagged strains were constructed as described (Snyder et al., 2019) and freshly prepared 10,000x stocks (500 mM) 3-indole acetic acid (IAA) (Sigma Aldrich) dissolved in 100% ethanol were used to treat yeast cultures (YPD) at 50 μM IAA. NatMX6 gene-replacement was performed by amplifying the *NatMX6* cassette from the p41Nat 1-F GW plasmid (Addgene #58546) using 45 bp primers flanking the *APS3 ORF*. Linear fragments were transformed into BY4742 and selected on YPD agar containing nourseothricin sulfate (100 μg/ml, GoldBio).

### Genome-wide cell death screen analysis

Screen analysis and batch correction were performed for the heat-ramp cell death assay previously reported Cell death was determined by quantifying microscopy colonies using a BioSpot reader at 18 h after heat-ramp (Teng et al., 2011; Teng et al., 2013). For this study, original images from four replicates plated at each of two dilutions were reacquired from the BioSpot reader and batch corrections were applied to systematically correct for undercounting at higher colony densities. Hit cutoff was 1.5 x IQR beyond 75^th^ percentile of all z-scores. Gene ontology (GO) component analysis was performed at the Saccharomyces Genome Database (SGD https://www.yeastgenome.org/).

### Heat-ramp cell death assays

Heat-ramp assays were performed on log phase cultures throughout this study as (30°C to 51°C, over 20 min, Figure 1A) (Teng et al., 2011; Teng and Hardwick, 2013), except the genome-wide screen performed on post-diauxic cells (linear 30°C to 62°C over 20 min) (Teng et al., 2013). Briefly, log-phase yeast grown on a roller drum at 30°C overnight (∼16 h), were normalized to 0.2 OD_600_/mL in 4-5 mL fresh YPD and grown for ∼3-4 h to log phase with adjustment to have all strains arrive at the same growth state approximately simultaneously. Equal cell numbers (100 μL of 0.4-0.5 OD_600_/mL YPD) were treated with a heat-ramp using an equilibrated programmable thermocycler. Heated and unheated samples (3 μl or 5 μl) were quickly spotted on agar plates in 5-fold serial dilutions in YPD, incubated 18-24 h at 30°C for semi-automated counting of microscopic colonies on a BioSpot Reader, or 2-days for colony visualization, and calculated as cfu/mL relative to untreated (**Supplementary Figure S5**).

*Cryptococcus* strains were grown in Sabouraud dextrose broth (SAB, BD Difco) for 2 days without shaking at 30°C. Cells were diluted 1:5 in fresh SAB and treated with a linear heat-ramp of 30°C to 52°C over ∼30min. Pre and post heat-ramp samples were spotted on SAB agar and incubated 48 h at 30°C before imaging.

### Immunoblot analysis

Approximately equal cell numbers (equivalent to 2.0 OD_600_) were pelleted and stored frozen at -80°C. For time-course experiments, cells were rapidly lysed with 192 μl 100% trichloroacetic acid (TCA), held on ice for at least 30 min, and cell pellets were washed three times with water, once with acetone, dried under vacuum for 10 min and stored frozen at -80°C (adapted from (Hughes Hallett et al., 2015). Frozen pellets from either protocol were resuspended in 200 μl sample buffer (62.5 mM Tris-HCl pH 6.8, 2% w/v SDS, 10% glycerol, 0.01% bromophenol blue, fresh 50 mM DTT, and Halt^TM^ protease inhibitor cocktail from ThermoFisher Scientific). Lysates were vortexed 4 x 45 sec with glass beads (Sigma Aldrich), maintained on ice between pulses, and heated to 95°C for 5 min before analysis by SDS-PAGE. Blots were probed with antibodies against flag-epitope (Sigma, 1:5000), PGK (Abcam, 1:5000), Pho8 (1:1000, Gregory Payne), Apm3 (1:5000, Sandra Lemmon), followed by HRP-conjugated anti-rabbit or anti-mouse (GE Healthcare, 1:5000).

### Cell death by vital dye staining

Following heat-ramp, yeast were stained with phloxine B (2 μg/mL, Fisher Scientific) or propidium iodide (10 μg/mL, Sigma Aldrich) and 10 μL stained cells loaded on a hemocytometer were imaged using a Zeiss AxioImager M2 (10x Olympus objective equipped with a Hamamatsu Orca R2 camera. Alternatively, 3 μL of stained cells were spotted directly onto YPD agar plates and imaged using an Applied Precision DeltaVision Elite microscrope system (GE Healthcare) equipped with 20xPh objective and automated stage in a 30°C incubator chamber. Time-lapse images of cells in fresh YPD with phloxine B (2 μg/mL, Fisher Scientific) were acquired every 10-20 min and analyzed using ImageJ/FIJI. Initial total cell count was calculated using phase images from the first imaging time point.

### Imaging vacuole membrane permeabilization

Yeast strains were grown overnight, normalized to equal cell numbers in fresh YPD (0.2 OD_600_/mL) and incubated ∼4 h. During the final 30 min, 1-mL aliquots of cell suspensions were stained at 30°C with FM4-64 (10 μg/mL, ThermoFisher Scientific) and/or CellTracker™ Blue CMAC (100 μM, 7-amino-4-chloromethylcoumarin, ThermoFisher Scientific). Stained cultures were re-normalized to equal cell numbers (0.5 OD_600_/mL), heat-ramp treated, and 50 μL were immobilized for 10 min on a concanavalin A-coated 8-well μ-Slide (Ibidi) containing 200 μL of fresh YPD, with addition of UVITEX 2B (1 μg/mL, Polysciences, Inc.) for cell wall staining. YPD was aspirated and replaced with 200 μL of the media type required for imaging. For cell death, fresh YPD contained phloxine B (2 μg/mL, Fisher Scientific). For imaging in FITC/GFP channels, YPD was replaced with 200 μL synthetic complete (SC) medium to avoid autofluorescence. Prototroph BY4709 strains were imaged in minimal SC^MIN^ (0.67% yeast nitrogen base without amino acids, 2% dextrose, 20 mg/L uracil), and BY4741/4742 strains were imaged in SC^AUX^ (Teng et al., 2018). Images were captured on a DeltaVision Elite microscope with a 60x (N.A. 1.42) oil immersion objective in a 30°C incubator chamber, and single image slices or 0.2 μm Z-stacks were deconvolved (Softworx, Applied Precision). To avoid phototoxicity, time-lapse microscopy used spaced time-lapse intervals (10 min), optical axis integration (OAI, z-sweep acquisition technology) and subsequent deconvolution. Images were analyzed with ImageJ/FIJI software.

## Acknowledgements

We thank Gregory Payne for Pho8/ALP antiserum, and Sandra Lemmon for Apm3 antiserum. We thank Leonid Kruglyak for assistance with plasmid “p41Nat 1-F GW”, Linda Huang for assistance with plasmid pFA6a-link-GFPEnvy-SpHis5, Pierre Morsomme for GFP-Sna2 plasmids, Susan Michaelis for the pRS303 plasmid, Jef Boeke for BY4709, and Scott Emr for SEY6210 strains. We thank Pierre Durand and Eugene Koonin for comments on this manuscript regarding the evolution of programmed cell death.

## SUPPLEMENTARY Movies

**Supplementary Movie S1. Movie for images in Figure 3C main text.**

**Supplementary Movie S2. Movie for images in Figure 6A main text.**

## SUPPLEMENTARY TABLES

**Supplementary Table S1. Genome-wide cell death screen results and Z-scores**

**Supplementary Table S2. Yeast strains used in each figure panel**

**Supplementary Table S3. Plasmid list and sequences of primers and gBlocks**

---

**Supplemental Figure S1.**
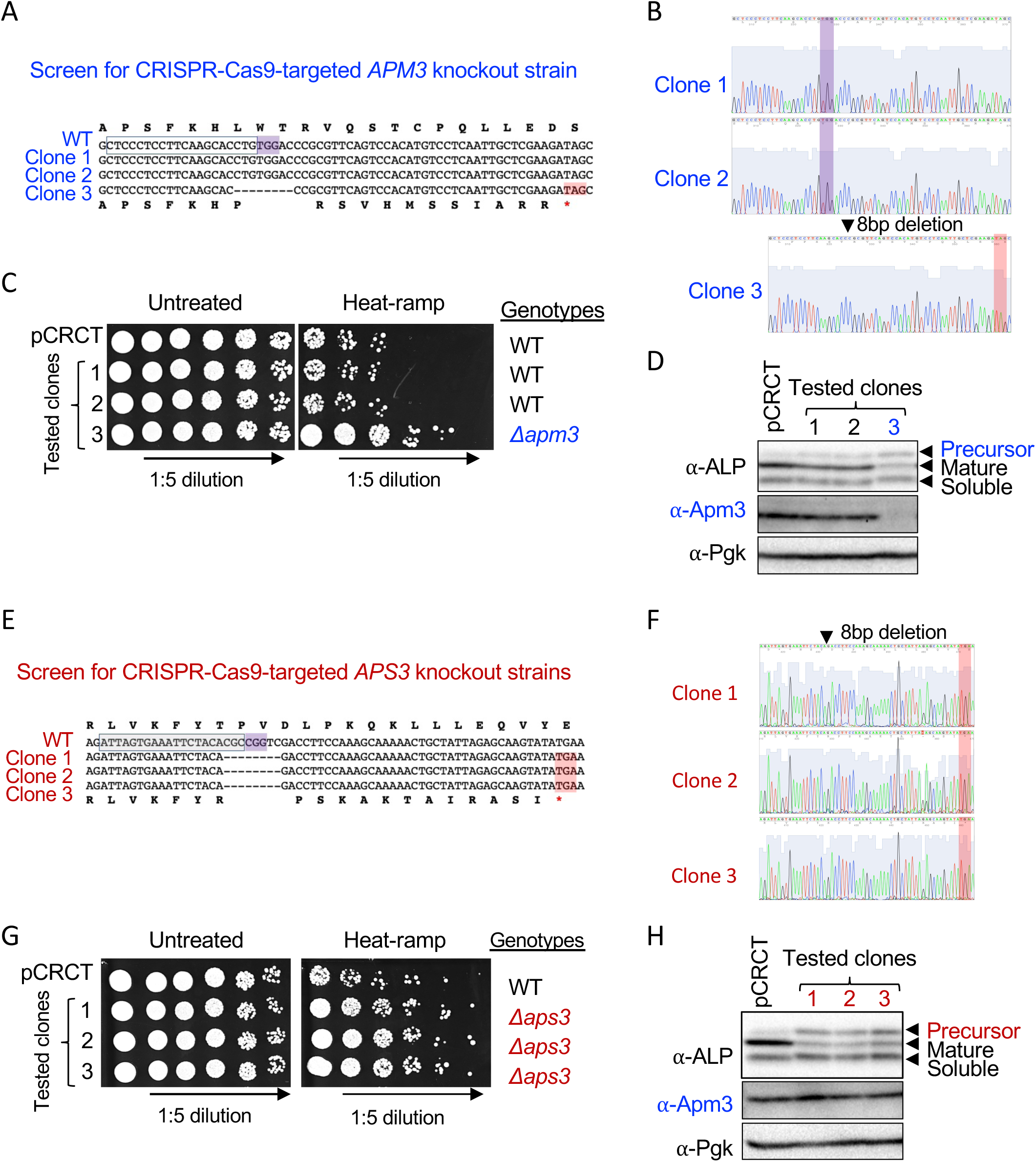
CRISPR-Cas9 design and sequencing for *APM3* and *APS3* knockout strains. [supports Figure 2 main text]. (A and E) Sequence alignments of genomic DNA from three colony-derived yeast clones (BY4709) transformed with gene-targeted or untargeted CRISPR pCRCT plasmids. CRISPR-Cas9 guide RNA sequence (box), PAM sequence (violet), and resulting in-frame stop codon (rose) following 8 bp deletion. (B and F) Sanger sequencing chromatograms for A and E. (C and G) Heat-ramp cell death assays for sequenced clones (30°C to 51°C as described in Figure 1A) confirm that death-resistance correlates with gene disruptions. (D and H) Tests for AP-3 trafficking function by assessing transport and processing of the precursor form of alkaline phosphatase (ALP/Pho8) to mature and soluble forms detected with anti-ALP antibody (from Gregory Payne, UCLA). Apm3 immunoblots confirm lack of expression in *Δapm3*, and evidence for AP-3 complex stability in *Δaps3*, assessed with anti-Apm3 antibodies provided by Dr. Sandra Lemmon, University of Miami (Panek et al., 1997).

**Supplementary Figure S2.**
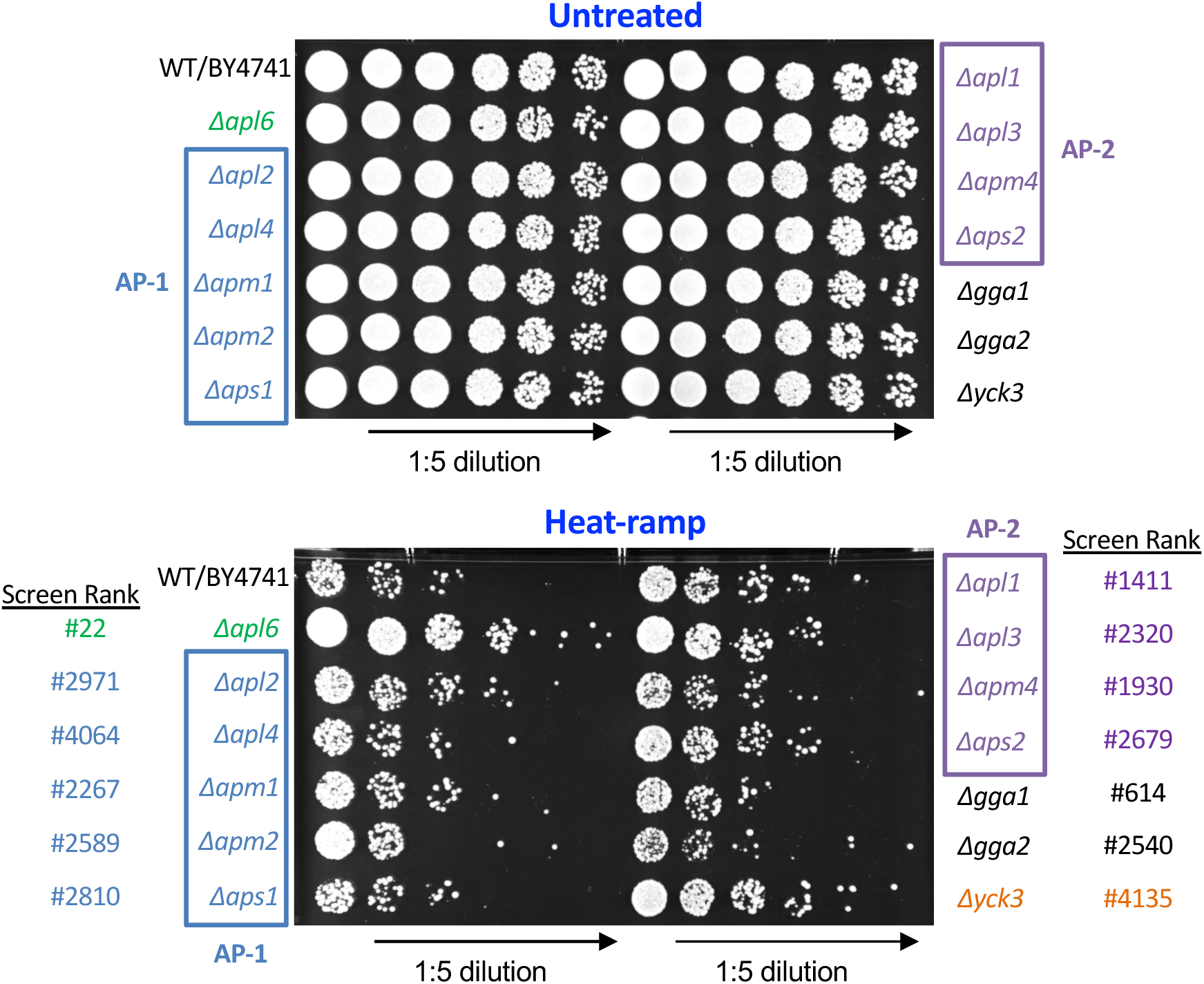
Cell death assay confirming lack of death-resistance for deletion strains of adaptor complexes AP-1 and AP-2. [supports Figure 2 main text]. Log phase yeast cultures of AP-1 (blue boxes), AP-2 (violet boxes) and other yeast knockout strains (BY4741 *MATa* YKO collection) were spotted on plates before (upper) and after (lower) heat-ramp treatment (20-min, 30°-51°C, from Figure 1A) and compared to the death-resistant AP-3 subunit Apl6 knockout (*Δapl6*). Ranks from the genome-wide heat-ramp screen in Figure 2A and Supplementary Table S1 are shown.

**Supplemental Figure S3.**
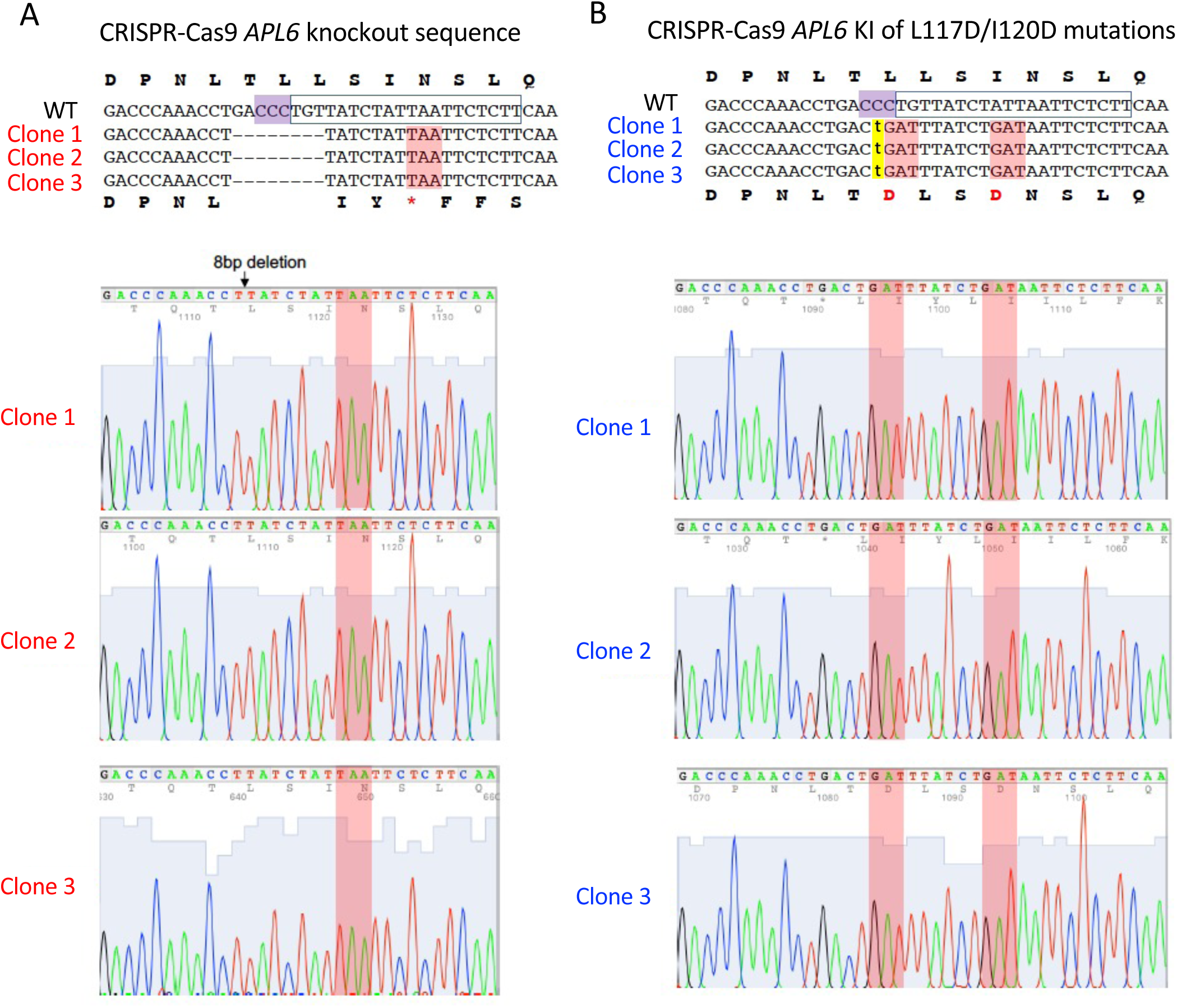
CRISPR-Cas9 design and sequencing for *APL6* knockin and knockout mutant strains. [supports Figure 4 main text]. (A) Genomic DNA sequence alignment of *APL6*-disrupted strains in BY4709 background, showing CRISPR-Cas9 guide RNA sequence (box), PAM sequence (violet), the resulting in-frame stop codon (rose) following 8 bp deletion, and corresponding Sanger sequencing chromatograms. (B) Genomic DNA sequence alignment of Arf1-interacting site mutations engineered into *APL6* knockin strains (BY4709). CRISPR-Cas9 guide RNA sequence (box); PAM sequence (violet); knockin Leu117Asp/Ile120Asp mutations (rose) based on Ren *et al*. Cell 2013; synonymous C>T nucleotide change engineered to eliminate PAM sequence (lower case, yellow highlight); corresponding Sanger sequencing chromatograms.

**Supplementary Figure S4.**
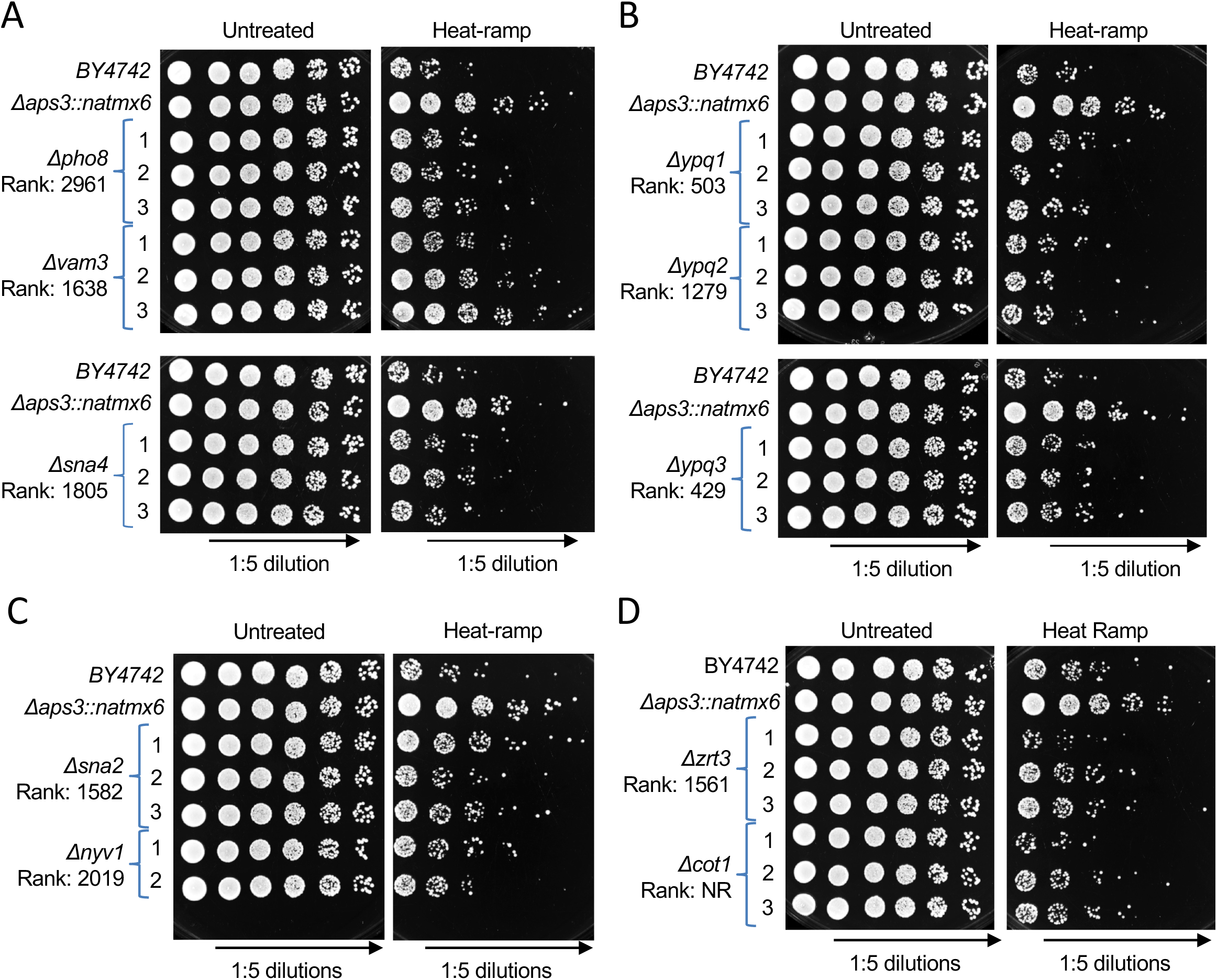
Retesting knockout strains of AP-3 cargo for cell death phenotypes. [supports Figure 4 main text]. Cell death assays for knockouts of AP-3 cargo proteins that contain a di-leucine motif (A and B), or a tyrosine motif (C), for interacting with AP-3, and other candidate cargo (D). Growth/survival of spotted yeast cultures before and after heat-ramp (20 min, 30℃ to 51℃, Figure 1A). Each strain from the BY4741 YKO collection was streaked onto rich (YPD) agar plates and three single cell-derived colonies (substrains #1-3) were tested in parallel with the AP-3 knockout *Δaps3* and its parental wild type (BY4742). Five-fold serial dilutions were plated (5 μL) and imaged after incubating 2-days at 30℃. Rank: z-scores for cell death resistance from the genome-wide screen.

**Supplementary Figure S5.**
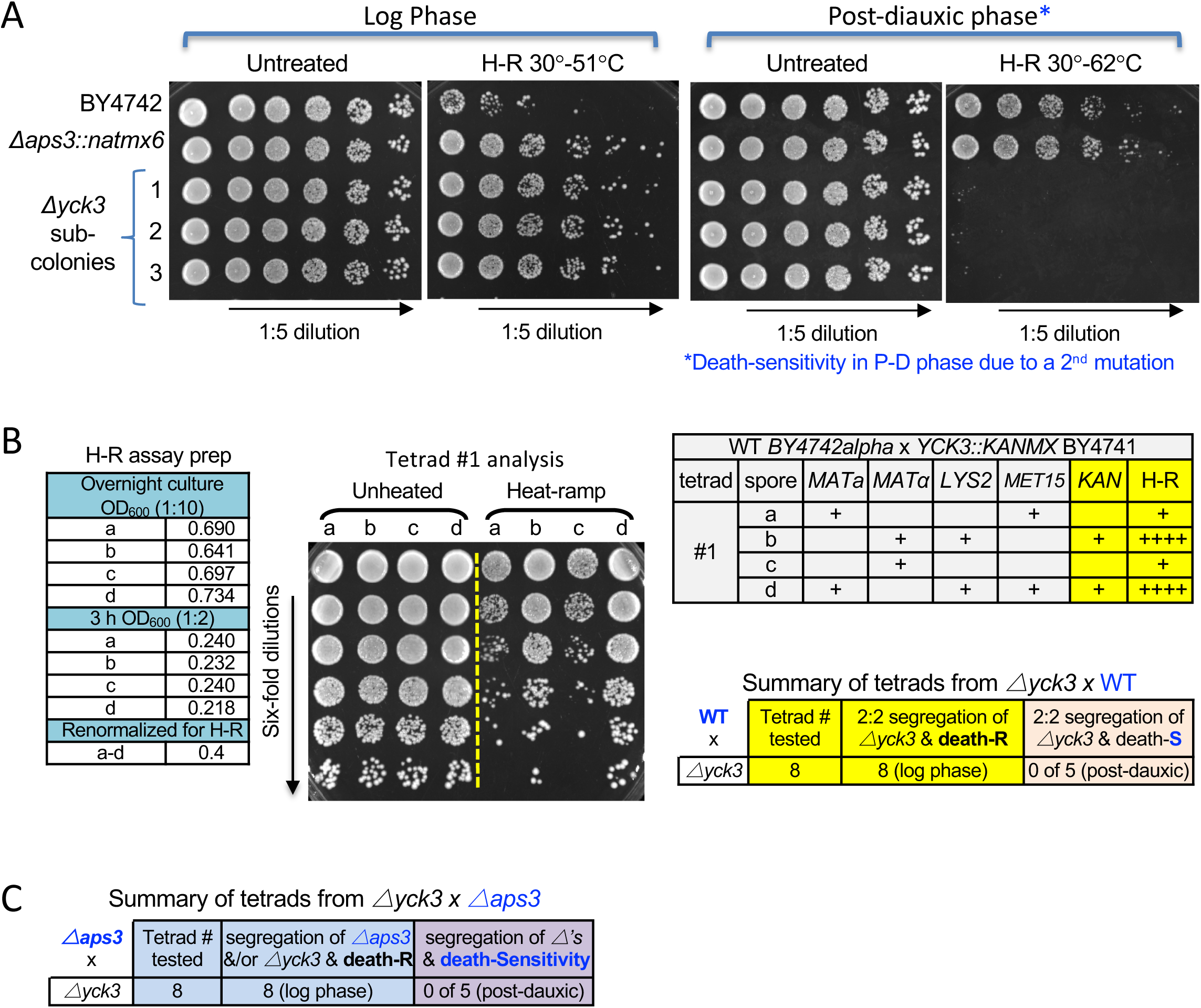
Tetrad analyses demonstrating deletion of *YCK3* confers death-resistance. [supports Figure 4 main text]. (A) Heat-ramp cell death assay for single cell-derived substrains of *Δyck3* (BY4741 YKO collection) and reference strains tested in log-phase (20-min, 30°C to 51°C, Figure 1A) and in post-dauxic phase used in the genome-wide screen (20-min, linear 30°C to 62°C). (B) Cell death assay results for example tetrad (4 spore-derived strains) from crossing WT x *Δyck3* and tested as in A. ODs of yeast cultures during sample preparation prior to heat-ramp (left) to minimize effects of metabolic state differences on cell death versus survival. Tetrad validation markers (upper right), reveal co-segregation of the knockout locus (*KanMX*) with death-resistance, summarized for all tetrads tested (lower right). Death-resistance (R), death-sensitive (S). (C) Summary of cell death results for all tetrads tested from crossing *Δyck3* x *Δaps3* in log and post-dauxic phases as in A. Cell death sensitivity in post-dauxic phase only is due to a secondary mutation that segregates independently, and is assumed to be present in the original *Δyck3* strain as it is present in crosses to both *Δaps3*, and to WT (B), while crosses *Δaps3* or other AP-3 knockouts to WT lack this mutation. <u>Heat-ramp method details</u>. Strains were streaked onto rich YPD agar, incubated 2-days at 30°C, and approximately matched numbers of cells from individual colonies (substrains) or the population, were grown in liquid YPD on a roller at 30°C for ∼16 h. Starting OD_600_ was recorded, cultures were normalized to 0.2 OD_600_/mL in 3 mL fresh YPD, and grown for 3 h with rotation at 30°C to reach log phase with closely matched OD_600_ between samples (∼0.45 OD_600_). Cultures were re-normalized to 0.4 OD_600_, 100 μL was transferred to 0.2 mL tubes for heat-ramp treatment in a thermocycler, and 3 μL of serial dilutions were immediately spotted on YPD agar and incubated 18-24h for BioSpot imaging, or 2-days for visual imaging on a ChemiDoc® Imager (Bio-Rad, CA, USA), exposure time 0.175 seconds.

**Supplementary Figure S6.**
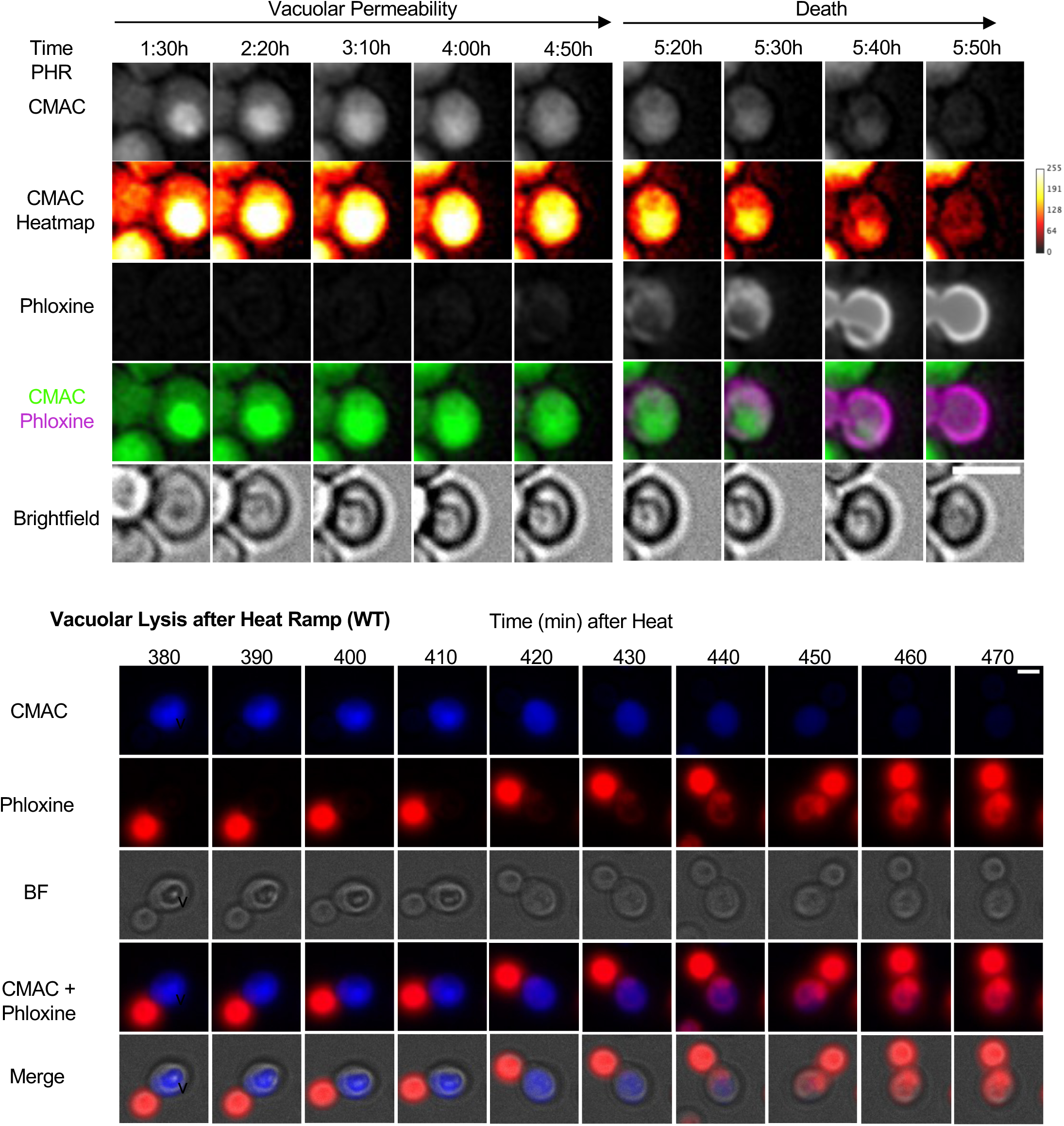
Vacuole membrane permeabilization after heat-ramp treatment. [supports Figure 6A main text]. Top: Another example as described in Figure 6A main test. Scale bar 2.5 μm. Bottom: Non-deconvolved micrographs of an independent experiment showing vacuole permeabilization and subsequent phloxine staining of WT cells after heat-ramp treatment. Scale bar 2.5 μm.

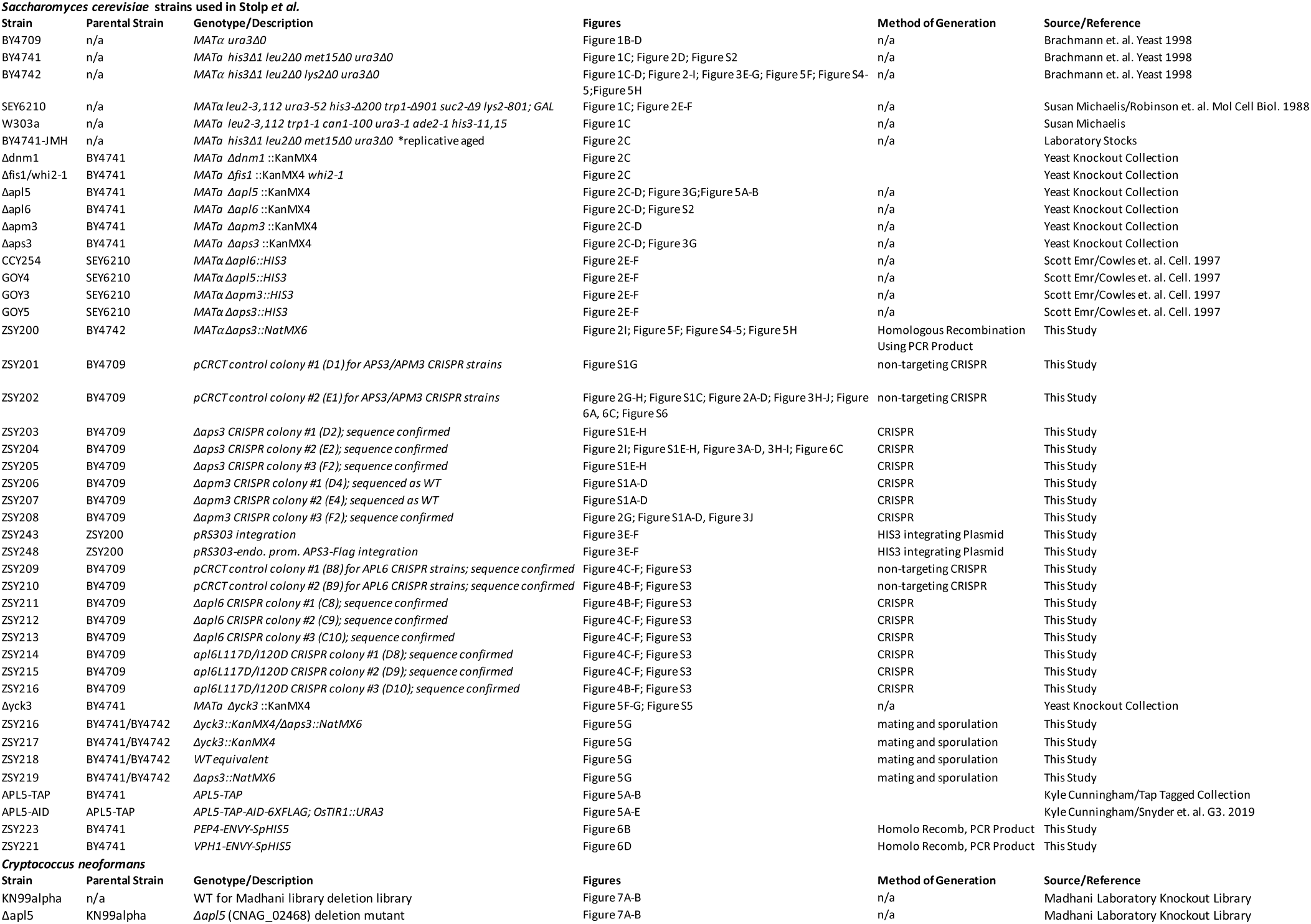

| **Plasmid** | **Description/Use** | **Source** |
| --- | --- | --- |
| p41Nat 1-F GW | natMX6 PCR template | Addgene 58546 |
| pCRCT | CRISPR Vector | Addgene 60621 |
| pFA6A-link-GFPEnvy-SpHis5 | PCR template for C' terminally tagging genes at endogenous locus | Addgene 60782 |
| pCRCT-APS3 disruption | insert 8 bp deletion within APS3 by CRISPR and HDR | This Study |
| pCRCT-APM3 disruption | insert 8 bp deletion within APM3 by CRISPR and HDR | This Study |
| pCRCT-APL6 disruption | insert 8 bp deletion within APL6 by CRISPR and HDR | This Study |
| pCRCT-APL6 L117D/I120D | insert substitution mutation in APL6 by CRISPR and HDR | This Study |
| pRS303 | HIS integrating vector | Susan Michaelis |
| ZDS100 | pRS303-*pAPS3* -APS3-FLAG | This Study |
| pRS416-*pSNA2* -GFP-SNA2 | GFP-SNA2 Expressing vector | Pierre Morsomme |

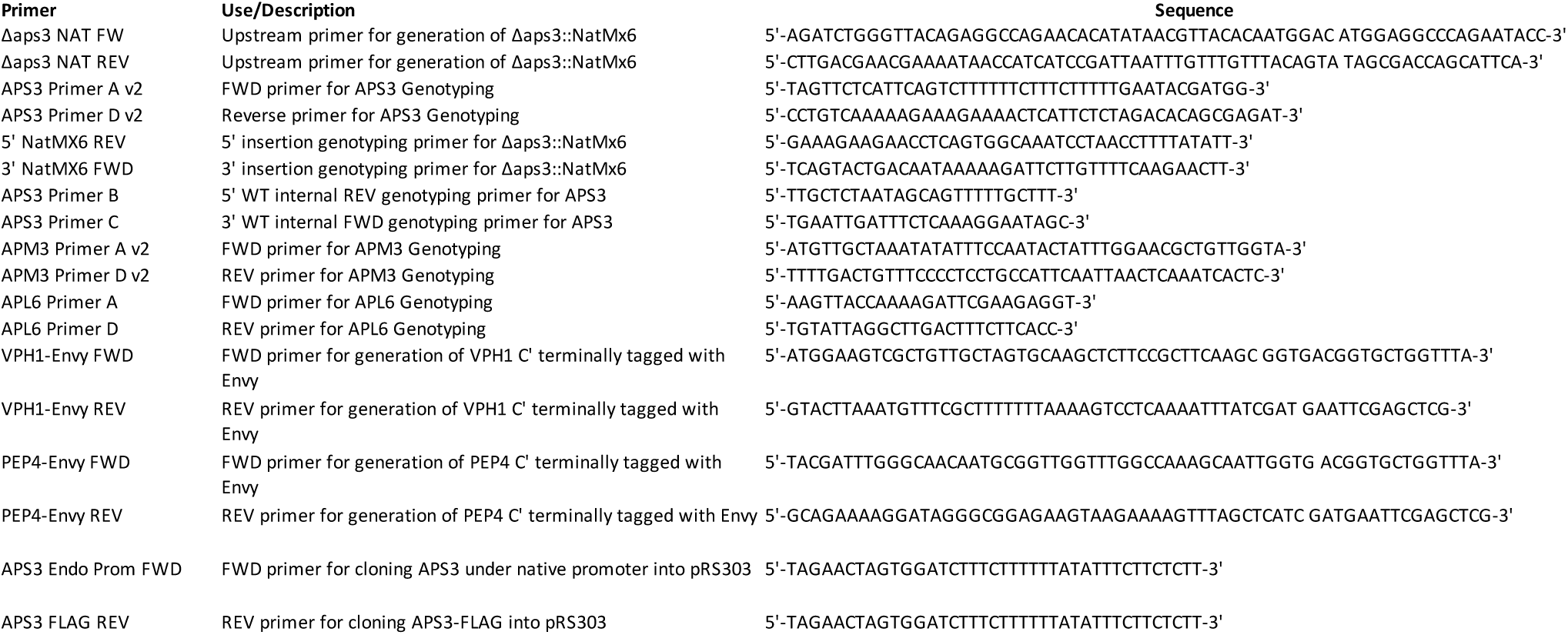

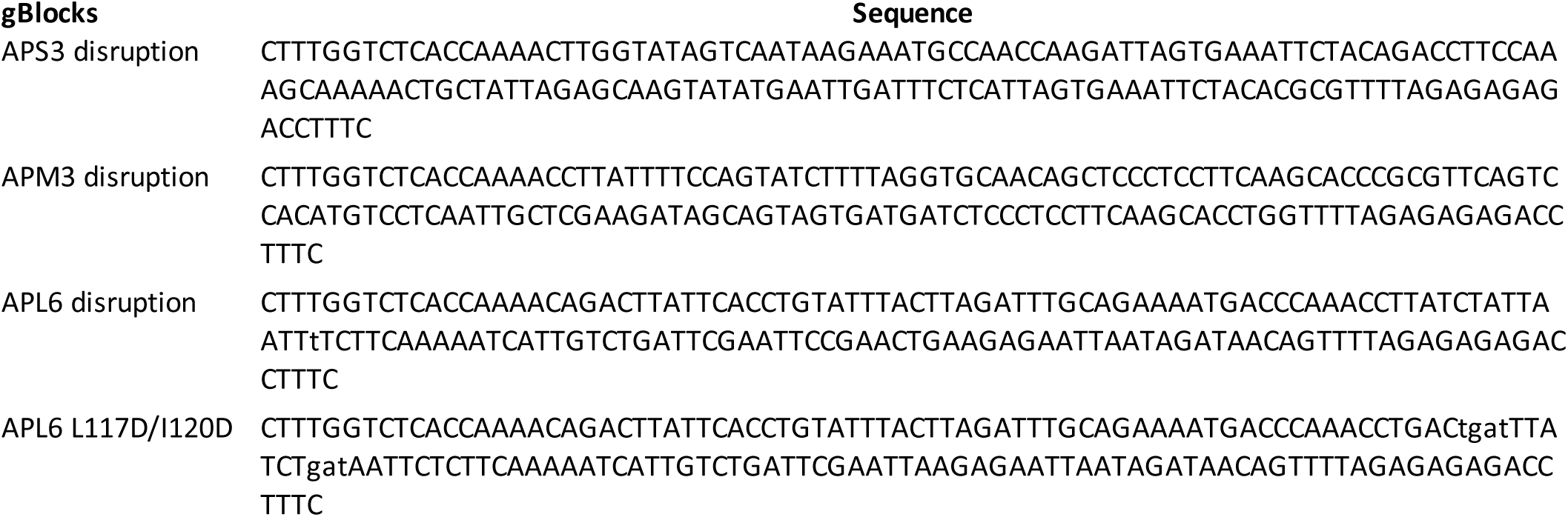

## REFERENCES

1. Ameisen, J.C. (2002). On the origin, evolution, and nature of programmed cell death: a timeline of four billion years. Cell Death Differ 9, 367–393.

2. Anand, V.C., Daboussi, L., Lorenz, T.C., and Payne, G.S. (2009). Genome-wide analysis of AP-3-dependent protein transport in yeast. Mol Biol Cell 20, 1592–1604.

3. Aouacheria, A., Cunningham, K.W., *Hardwick, J.M., Palkova, Z., Powers, T., Severin, F.F., and Vachova, L. (2018). Comment on “Sterilizing immunity in the lung relies on targeting fungal apoptosis-like programmed cell death”. Science 360, *Corresponding author.

4. Bao, Z., Xiao, H., Liang, J., Zhang, L., Xiong, X., Sun, N., Si, T., and Zhao, H. (2015). Homology-integrated CRISPR-Cas (HI-CRISPR) system for one-step multigene disruption in Saccharomyces cerevisiae. ACS Synth Biol 4, 585–594.

5. Baruffini, E., Ruotolo, R., Bisceglie, F., Montalbano, S., Ottonello, S., Pelosi, G., Buschini, A., and Lodi, T. (2020). Mechanistic insights on the mode of action of an antiproliferative thiosemicarbazone-nickel complex revealed by an integrated chemogenomic profiling study. Sci Rep 10, 10524.

6. Brett, C.L., Plemel, R.L., Lobingier, B.T., Vignali, M., Fields, S., and Merz, A.J. (2008). Efficient termination of vacuolar Rab GTPase signaling requires coordinated action by a GAP and a protein kinase. J Cell Biol 182, 1141–1151.

7. Buelto, D., Hung, C.W., Aoh, Q.L., Lahiri, S., and Duncan, M.C. (2020). Plasma membrane to vacuole traffic induced by glucose starvation requires Gga2-dependent sorting at the trans-Golgi network. Biol Cell 112, 349–367.

8. Cabrera, M., Langemeyer, L., Mari, M., Rethmeier, R., Orban, I., Perz, A., Brocker, C., Griffith, J., Klose, D., Steinhoff, H.J., et al. (2010). Phosphorylation of a membrane curvature-sensing motif switches function of the HOPS subunit Vps41 in membrane tethering. J Cell Biol 191, 845–859.

9. Cabrera, M., Ostrowicz, C.W., Mari, M., LaGrassa, T.J., Reggiori, F., and Ungermann, C. (2009). Vps41 phosphorylation and the Rab Ypt7 control the targeting of the HOPS complex to endosome-vacuole fusion sites. Mol Biol Cell 20, 1937–1948.

10. Casler, J.C., and Glick, B.S. (2020). A microscopy-based kinetic analysis of yeast vacuolar protein sorting. Elife 9.

11. Chaves, S.R., Rego, A., Martins, V.M., Santos-Pereira, C., Sousa, M.J., and Côrte-Real, M. (2021). Regulation of Cell Death Induced by Acetic Acid in Yeasts. Frontiers in Cell and Developmental Biology 9.

12. Cheng, W.C., Teng, X., Park, H.K., Tucker, C.M., Dunham, M.J., and Hardwick, J.M. (2008). Fis1 deficiency selects for compensatory mutations responsible for cell death and growth control defects. Cell Death Differ 15, 1838–1846.

13. Clavé, C., Dyrka, W., Granger-Farbos, A., Pinson, B., Saupe, S.J., and Daskalov, A. (2021). Fungal gasdermin-like proteins are controlled by proteolytic cleavage. bioRxiv, 2021.2006.2003.446900.

14. Cowles, C.R., Odorizzi, G., Payne, G.S., and Emr, S.D. (1997a). The AP-3 adaptor complex is essential for cargo-selective transport to the yeast vacuole. Cell 91, 109–118.

15. Cowles, C.R., Snyder, W.B., Burd, C.G., and Emr, S.D. (1997b). Novel Golgi to vacuole delivery pathway in yeast: identification of a sorting determinant and required transport component. EMBO J 16, 2769–2782.

16. Daboussi, L., Costaguta, G., and Payne, G.S. (2012). Phosphoinositide-mediated clathrin adaptor progression at the trans-Golgi network. Nature Cell Biology 14, 239–248.

17. Darsow, T., Burd, C.G., and Emr, S.D. (1998). Acidic di-leucine motif essential for AP-3-dependent sorting and restriction of the functional specificity of the Vam3p vacuolar t-SNARE. J Cell Biol 142, 913–922.

18. Dell’Angelica, E.C., and Bonifacino, J.S. (2019). Coatopathies: Genetic Disorders of Protein Coats. Annu Rev Cell Dev Biol 35, 131–168.

19. Dong, Y., Hu, J., Fan, L., and Chen, Q. (2017). RNA-Seq-based transcriptomic and metabolomic analysis reveal stress responses and programmed cell death induced by acetic acid in Saccharomyces cerevisiae. Sci Rep 7, 42659.

20. Durand, P.M.C. (2020). Programmed cell death at the levels of selection. In The Evolutionary Origins of Life and Death (Chicago, Illinois: The University of Chicago Press).

21. Eastwood, M.D., Cheung, S.W., Lee, K.Y., Moffat, J., and Meneghini, M.D. (2012). Developmentally programmed nuclear destruction during yeast gametogenesis. Dev Cell 23, 35–44.

22. Eastwood, M.D., Cheung, S.W., and Meneghini, M.D. (2013). Programmed nuclear destruction in yeast: self-eating by vacuolar lysis. Autophagy 9, 263–265.

23. Eastwood, M.D., and Meneghini, M.D. (2015). Developmental Coordination of Gamete Differentiation with Programmed Cell Death in Sporulating Yeast. Eukaryot Cell 14, 858–867.

24. Erez, Z., Steinberger-Levy, I., Shamir, M., Doron, S., Stokar-Avihail, A., Peleg, Y., Melamed, S., Leavitt, A., Savidor, A., Albeck, S., et al. (2017). Communication between viruses guides lysis-lysogeny decisions. Nature 541, 488–493.

25. Fannjiang, Y., Cheng, W.C., Lee, S.J., Qi, B., Pevsner, J., McCaffery, J.M., Hill, R.B., Basanez, G., and Hardwick, J.M. (2004). Mitochondrial fission proteins regulate programmed cell death in yeast. Genes Dev 18, 2785–2797.

26. Galluzzi, L., Vitale, I., Aaronson, S.A., Abrams, J.M., Adam, D., Agostinis, P., Alnemri, E.S., Altucci, L., Amelio, I., Andrews, D.W., et al. (2018). Molecular mechanisms of cell death: recommendations of the Nomenclature Committee on Cell Death 2018. Cell Death Differ 25, 486–541.

27. Gao, J., Chau, S., Chowdhury, F., Zhou, T., Hossain, S., McQuibban, G.A., and Meneghini, M.D. (2019). Meiotic viral attenuation through an ancestral apoptotic pathway. Proceedings of the National Academy of Sciences 116, 16454–16462.

28. Gietz, R.D., and Schiestl, R.H. (2007). High-efficiency yeast transformation using the LiAc/SS carrier DNA/PEG method. Nat Protoc 2, 31–34.

29. Goncalves, A.P., Heller, J., Daskalov, A., Videira, A., and Glass, N.L. (2017). Regulated Forms of Cell Death in Fungi. Front Microbiol 8, 1837.

30. Heller, J., Clave, C., Gladieux, P., Saupe, S.J., and Glass, N.L. (2018). NLR surveillance of essential SEC-9 SNARE proteins induces programmed cell death upon allorecognition in filamentous fungi. Proc Natl Acad Sci U S A 115, E2292–E2301.

31. Hirst, J., Lindsay, M.R., and Robinson, M.S. (2001). GGAs: roles of the different domains and comparison with AP-1 and clathrin. Mol Biol Cell 12, 3573–3588.

32. Hughes Hallett, J.E., Luo, X., and Capaldi, A.P. (2015). Snf1/AMPK promotes the formation of Kog1/Raptor-bodies to increase the activation threshold of TORC1 in budding yeast. Elife 4.

33. Iranzo, J., Lobkovsky, A.E., Wolf, Y.I., and Koonin, E.V. (2014). Virus-host arms race at the joint origin of multicellularity and programmed cell death. Cell Cycle 13, 3083–3088.

34. Ivanovska, I., and Hardwick, J.M. (2005). Viruses activate a genetically conserved cell death pathway in a unicellular organism. J Cell Biol 170, 391–399.

35. Jarolim, S., Ayer, A., Pillay, B., Gee, A.C., Phrakaysone, A., Perrone, G.G., Breitenbach, M., and Dawes, I.W. (2013). Saccharomyces cerevisiae genes involved in survival of heat shock. G3 (Bethesda) 3, 2321–2333.

36. Johnson, A.G., Wein, T., Mayer, M.L., Duncan-Lowey, B., Yirmiya, E., Oppenheimer-Shaanan, Y., Amitai, G., Sorek, R., and Kranzusch, P.J. (2021). Bacterial gasdermins reveal an ancient mechanism of cell death. bioRxiv, 2021.2006.2007.447441.

37. Karim, M.A., McNally, E.K., Samyn, D.R., Mattie, S., and Brett, C.L. (2018). Rab-Effector-Kinase Interplay Modulates Intralumenal Fragment Formation during Vacuole Fusion. Dev Cell 47, 80–97 e86.

38. Kayagaki, N., Kornfeld, O.S., Lee, B.L., Stowe, I.B., O’Rourke, K., Li, Q., Sandoval, W., Yan, D., Kang, J., Xu, M., et al. (2021). NINJ1 mediates plasma membrane rupture during lytic cell death. Nature 591, 131–136.

39. Kim, A., and Cunningham, K.W. (2015). A LAPF/phafin1-like protein regulates TORC1 and lysosomal membrane permeabilization in response to endoplasmic reticulum membrane stress. Mol Biol Cell 26, 4631–4645.

40. Kim, H., Kim, A., and Cunningham, K.W. (2012). Vacuolar H+-ATPase (V-ATPase) promotes vacuolar membrane permeabilization and nonapoptotic death in stressed yeast. J Biol Chem 287, 19029–19039.

41. Klionsky, D.J., and Emr, S.D. (1989). Membrane protein sorting: biosynthesis, transport and processing of yeast vacuolar alkaline phosphatase. EMBO J 8, 2241–2250.

42. Klionsky, D.J., and Eskelinen, E.L. (2014). The vacuole versus the lysosome: when size matters. Autophagy 10, 185–187.

43. Koonin, E.V., and Zhang, F. (2017). Coupling immunity and programmed cell suicide in prokaryotes: Life-or-death choices. Bioessays 39, 1–9.

44. Kuida, K., Zheng, T.S., Na, S., Kuan, C., Yang, D., Karasuyama, H., Rakic, P., and Flavell, R.A. (1996). Decreased apoptosis in the brain and premature lethality in CPP32-deficient mice. Nature 384, 368–372.

45. Kulkarni, M., Stolp, Z.D., and Hardwick, J.M. (2019). Targeting intrinsic cell death pathways to control fungal pathogens. Biochem Pharmacol.

46. Kwolek-Mirek, M., and Zadrag-Tecza, R. (2014). Comparison of methods used for assessing the viability and vitality of yeast cells. FEMS Yeast Res 14, 1068–1079.

47. LaGrassa, T.J., and Ungermann, C. (2005). The vacuolar kinase Yck3 maintains organelle fragmentation by regulating the HOPS tethering complex. J Cell Biol 168, 401–414.

48. Lapinskas, P.J., Cunningham, K.W., Liu, X.F., Fink, G.R., and Culotta, V.C. (1995). Mutations in PMR1 suppress oxidative damage in yeast cells lacking superoxide dismutase. Mol Cell Biol 15, 1382–1388.

49. Lawrence, G., Brown, C.C., Flood, B.A., Karunakaran, S., Cabrera, M., Nordmann, M., Ungermann, C., and Fratti, R.A. (2014). Dynamic association of the PI3P-interacting Mon1-Ccz1 GEF with vacuoles is controlled through its phosphorylation by the type 1 casein kinase Yck3. Mol Biol Cell 25, 1608–1619.

50. Lemasters, J.J., Theruvath, T.P., Zhong, Z., and Nieminen, A.L. (2009). Mitochondrial calcium and the permeability transition in cell death. Biochim Biophys Acta 1787, 1395–1401.

51. Leuenberger, P., Ganscha, S., Kahraman, A., Cappelletti, V., Boersema, P.J., von Mering, C., Claassen, M., and Picotti, P. (2017). Cell-wide analysis of protein thermal unfolding reveals determinants of thermostability. Science 355.

52. Levi, S.K., Bhattacharyya, D., Strack, R.L., Austin, J.R., 2nd, and Glick, B.S. (2010). The yeast GRASP Grh1 colocalizes with COPII and is dispensable for organizing the secretory pathway. Traffic 11, 1168–1179.

53. Lindsten, T., Golden, J.A., Zong, W.X., Minarcik, J., Harris, M.H., and Thompson, C.B. (2003). The proapoptotic activities of Bax and Bak limit the size of the neural stem cell pool. J Neurosci 23, 11112–11119.

54. Liu, X., Zhang, Z., Ruan, J., Pan, Y., Magupalli, V.G., Wu, H., and Lieberman, J. (2016). Inflammasome-activated gasdermin D causes pyroptosis by forming membrane pores. Nature 535, 153–158.

55. Llinares, E., Barry, A.O., and Andre, B. (2015). The AP-3 adaptor complex mediates sorting of yeast and mammalian PQ-loop-family basic amino acid transporters to the vacuolar/lysosomal membrane. Sci Rep 5, 16665.

56. Lockshin, R.A., and Williams, C.M. (1965). Programmed Cell Death–I. Cytology of Degeneration in the Intersegmental Muscles of the Pernyi Silkmoth. J Insect Physiol 11, 123–133.

57. Manandhar, S.P., Siddiqah, I.M., Cocca, S.M., and Gharakhanian, E. (2020). A kinase cascade on the yeast lysosomal vacuole regulates its membrane dynamics: conserved kinase Env7 is phosphorylated by casein kinase Yck3. J Biol Chem 295, 12262–12278.

58. McNally, E.K., Karim, M.A., and Brett, C.L. (2017). Selective Lysosomal Transporter Degradation by Organelle Membrane Fusion. Dev Cell 40, 151–167.

59. Minina, E.A., Staal, J., Alvarez, V.E., Berges, J.A., Berman-Frank, I., Beyaert, R., Bidle, K.D., Bornancin, F., Casanova, M., Cazzulo, J.J., et al. (2020). Classification and Nomenclature of Metacaspases and Paracaspases: No More Confusion with Caspases. Mol Cell 77, 927–929.

60. Minois, N., Frajnt, M., Wilson, C., and Vaupel, J.W. (2005). Advances in measuring lifespan in the yeast Saccharomyces cerevisiae. Proc Natl Acad Sci U S A 102, 402–406.

61. Morawska, M., and Ulrich, H.D. (2013). An expanded tool kit for the auxin-inducible degron system in budding yeast. Yeast 30, 341–351.

62. Morris, K.L., Buffalo, C.Z., Sturzel, C.M., Heusinger, E., Kirchhoff, F., Ren, X., and Hurley, J.H. (2018). HIV-1 Nefs Are Cargo-Sensitive AP-1 Trimerization Switches in Tetherin Downregulation. Cell 174, 659–671 e614.

63. Nagata, S., and Segawa, K. (2021). Sensing and clearance of apoptotic cells. Curr Opin Immunol 68, 1–8.

64. Nie, Z., Boehm, M., Boja, E.S., Vass, W.C., Bonifacino, J.S., Fales, H.M., and Randazzo, P.A. (2003). Specific regulation of the adaptor protein complex AP-3 by the Arf GAP AGAP1. Dev Cell 5, 513–521.

65. Nishimura, K., and Kanemaki, M.T. (2014). Rapid Depletion of Budding Yeast Proteins via the Fusion of an Auxin-Inducible Degron (AID). Curr Protoc Cell Biol 64, 20 29 21-16.

66. Odorizzi, G., Cowles, C.R., and Emr, S.D. (1998). The AP-3 complex: a coat of many colours. Trends Cell Biol 8, 282–288.

67. Ooi, C.E., Dell’Angelica, E.C., and Bonifacino, J.S. (1998). ADP-Ribosylation factor 1 (ARF1) regulates recruitment of the AP-3 adaptor complex to membranes. J Cell Biol 142, 391–402.

68. Panek, H.R., Stepp, J.D., Engle, H.M., Marks, K.M., Tan, P.K., Lemmon, S.K., and Robinson, L.C. (1997). Suppressors of YCK-encoded yeast casein kinase 1 deficiency define the four subunits of a novel clathrin AP-like complex. EMBO J 16, 4194–4204.

69. Pokrzywa, W., Guerriat, B., Dodzian, J., and Morsomme, P. (2009). Dual sorting of the Saccharomyces cerevisiae vacuolar protein Sna4p. Eukaryot Cell 8, 278–286.

70. Ramisetty, B.C.M., Natarajan, B., and Santhosh, R.S. (2015). mazEF-mediated programmed cell death in bacteria: “What is this?”. Critical Reviews in Microbiology 41, 89–100.

71. Ren, X., Farias, G.G., Canagarajah, B.J., Bonifacino, J.S., and Hurley, J.H. (2013). Structural basis for recruitment and activation of the AP-1 clathrin adaptor complex by Arf1. Cell 152, 755–767.

72. Renard, H.F., Demaegd, D., Guerriat, B., and Morsomme, P. (2010). Efficient ER exit and vacuole targeting of yeast Sna2p require two tyrosine-based sorting motifs. Traffic 11, 931–946.

73. Robert, V.A., and Casadevall, A. (2009). Vertebrate endothermy restricts most fungi as potential pathogens. J Infect Dis 200, 1623–1626.

74. Roberts, A.W., Davids, M.S., Pagel, J.M., Kahl, B.S., Puvvada, S.D., Gerecitano, J.F., Kipps, T.J., Anderson, M.A., Brown, J.R., Gressick, L., et al. (2016). Targeting BCL2 with Venetoclax in Relapsed Chronic Lymphocytic Leukemia. N Engl J Med 374, 311–322.

75. Robinson, M.S., and Bonifacino, J.S. (2001). Adaptor-related proteins. Curr Opin Cell Biol 13, 444–453.

76. Ruan, J., Xia, S., Liu, X., Lieberman, J., and Wu, H. (2018). Cryo-EM structure of the gasdermin A3 membrane pore. Nature 557, 62–67.

77. Schoppe, J., Mari, M., Yavavli, E., Auffarth, K., Cabrera, M., Walter, S., Frohlich, F., and Ungermann, C. (2020). AP-3 vesicle uncoating occurs after HOPS-dependent vacuole tethering. EMBO J 39, e105117.

78. Seaman, M.N., Sowerby, P.J., and Robinson, M.S. (1996). Cytosolic and membrane-associated proteins involved in the recruitment of AP-1 adaptors onto the trans-Golgi network. J Biol Chem 271, 25446–25451.

79. Segarra, V.A., Boettner, D.R., and Lemmon, S.K. (2015). Atg27 tyrosine sorting motif is important for its trafficking and Atg9 localization. Traffic 16, 365–378.

80. Simpson, F., Peden, A.A., Christopoulou, L., and Robinson, M.S. (1997). Characterization of the adaptor-related protein complex, AP-3. J Cell Biol 137, 835–845.

81. Slubowski, C.J., Funk, A.D., Roesner, J.M., Paulissen, S.M., and Huang, L.S. (2015). Plasmids for C-terminal tagging in Saccharomyces cerevisiae that contain improved GFP proteins, Envy and Ivy. Yeast 32, 379–387.

82. Smith, R.P., Barraza, I., Quinn, R.J., and Fortoul, M.C. (2020). The mechanisms and cell signaling pathways of programmed cell death in the bacterial world. Int Rev Cell Mol Biol 352, 1–53.

83. Snyder, N.A., Kim, A., Kester, L., Gale, A.N., Studer, C., Hoepfner, D., Roggo, S., Helliwell, S.B., and Cunningham, K.W. (2019). Auxin-Inducible Depletion of the Essentialome Suggests Inhibition of TORC1 by Auxins and Inhibition of Vrg4 by SDZ 90-215, a Natural Antifungal Cyclopeptide. G3 (Bethesda) 9, 829–840.

84. Sousa, M., Duarte, A.M., Fernandes, T.R., Chaves, S.R., Pacheco, A., Leao, C., Corte-Real, M., and Sousa, M.J. (2013). Genome-wide identification of genes involved in the positive and negative regulation of acetic acid-induced programmed cell death in Saccharomyces cerevisiae. BMC Genomics 14, 838. https://doi.org/810.1186/1471-2164-1114-1838.

85. Spencer, S.L., Gaudet, S., Albeck, J.G., Burke, J.M., and Sorger, P.K. (2009). Non-genetic origins of cell-to-cell variability in TRAIL-induced apoptosis. Nature 459, 428–432.

86. Stepp, J.D., Huang, K., and Lemmon, S.K. (1997). The yeast adaptor protein complex, AP-3, is essential for the efficient delivery of alkaline phosphatase by the alternate pathway to the vacuole. J Cell Biol 139, 1761–1774.

87. Stott, K.E., Loyse, A., Jarvis, J.N., Alufandika, M., Harrison, T.S., Mwandumba, H.C., Day, J.N., Lalloo, D.G., Bicanic, T., Perfect, J.R., et al. (2021). Cryptococcal meningoencephalitis: time for action. The Lancet Infectious Diseases.

88. Sun, B., Chen, L., Cao, W., Roth, A.F., and Davis, N.G. (2004). The yeast casein kinase Yck3p is palmitoylated, then sorted to the vacuolar membrane with AP-3-dependent recognition of a YXXPhi adaptin sorting signal. Mol Biol Cell 15, 1397–1406.

89. Teng, X., Cheng, W.C., Qi, B., Yu, T.X., Ramachandran, K., Boersma, M.D., Hattier, T., Lehmann, P.V., Pineda, F.J., and Hardwick, J.M. (2011). Gene-dependent cell death in yeast. Cell Death Dis 2, e188. DOI: 110.1038/cddis.2011.1072.

90. Teng, X., Dayhoff-Brannigan, M., Cheng, W.C., Gilbert, C.E., Sing, C.N., Diny, N.L., Wheelan, S.J., Dunham, M.J., Boeke, J.D., Pineda, F.J., et al. (2013). Genome-wide consequences of deleting any single gene. Mol Cell 52, 485–494.

91. Teng, X., and Hardwick, J.M. (2013). Quantification of genetically controlled cell death in budding yeast. Methods Mol Biol 1004, 161–170.

92. Teng, X., and Hardwick, J.M. (2015). Cell death in genome evolution. Semin Cell Dev Biol 39, 3–11.

93. Teng, X., Yau, E., Sing, C., and Hardwick, J.M. (2018). Whi2 signals low leucine availability to halt yeast growth and cell death. FEMS Yeast Res 18, https://doi.org/10.1093/femsyr/foy1095.

94. Todd, R.T., and Selmecki, A. (2020). Expandable and reversible copy number amplification drives rapid adaptation to antifungal drugs. Elife 9.

95. Velazquez, R., Zamora, E., Alvarez, M., Alvarez, M.L., and Ramirez, M. (2016). Using mixed inocula of Saccharomyces cerevisiae killer strains to improve the quality of traditional sparkling-wine. Food Microbiol 59, 150–160.

96. Vilela-Moura, A., Schuller, D., Mendes-Faia, A., Silva, R.D., Chaves, S.R., Sousa, M.J., and Corte-Real, M. (2011). The impact of acetate metabolism on yeast fermentative performance and wine quality: reduction of volatile acidity of grape musts and wines. Appl Microbiol Biotechnol 89, 271–280.

97. Vowels, J.J., and Payne, G.S. (1998). A dileucine-like sorting signal directs transport into an AP-3-dependent, clathrin-independent pathway to the yeast vacuole. EMBO J 17, 2482–2493.

98. Watson, C.J., and Khaled, W.T. (2020). Mammary development in the embryo and adult: new insights into the journey of morphogenesis and commitment. Development 147.

99. Yang, X., Zhang, W., Wen, X., Bulinski, P.J., Chomchai, D.A., Arines, F.M., Liu, Y.-Y., Sprenger, S., Teis, D., Klionsky, D.J., et al. (2020). TORC1 regulates vacuole membrane composition through ubiquitin- and ESCRT-dependent microautophagy. Journal of Cell Biology 219.

100. Yesbolatova, A., Saito, Y., Kitamoto, N., Makino-Itou, H., Ajima, R., Nakano, R., Nakaoka, H., Fukui, K., Gamo, K., Tominari, Y., et al. (2020). The auxin-inducible degron 2 technology provides sharp degradation control in yeast, mammalian cells, and mice. Nat Commun 11, 5701.

